# The microbiome diversifies *N*-acyl lipid pools - including short-chain fatty acid-derived compounds

**DOI:** 10.1101/2024.10.31.621412

**Authors:** Helena Mannochio-Russo, Vincent Charron-Lamoureux, Martijn van Faassen, Santosh Lamichhane, Wilhan D. Gonçalves Nunes, Victoria Deleray, Abubaker Patan, Kyle Vittali, Prajit Rajkumar, Yasin El Abiead, Haoqi Nina Zhao, Paulo Wender Portal Gomes, Ipsita Mohanty, Carlynda Lee, Aidan Sund, Meera Sharma, Yuanhao Liu, David Pattynama, Gregory T. Walker, Grant J. Norton, Lora Khatib, Mohammadsobhan S. Andalibi, Crystal X. Wang, Ronald J. Ellis, David J. Moore, Jennifer E. Iudicello, Donald Franklin, Scott Letendre, Loryn Chin, Corinn Walker, Simone Renwick, Jasmine Zemlin, Michael J. Meehan, Xinyang Song, Dennis Kasper, Zachary Burcham, Jane J. Kim, Sejal Kadakia, Manuela Raffatellu, Lars Bode, Karsten Zengler, Mingxun Wang, Dionicio Siegel, Rob Knight, Pieter C. Dorrestein

## Abstract

*N*-acyl lipids are important mediators of several biological processes including immune function and stress response. To enhance the detection of *N*-acyl lipids with untargeted mass spectrometry-based metabolomics, we created a reference spectral library retrieving *N*-acyl lipid patterns from 2,700 public datasets, identifying 851 *N*-acyl lipids that were detected 356,542 times. 777 are not documented in lipid structural databases, with 18% of these derived from short-chain fatty acids and found in the digestive tract and other organs. Their levels varied with diet, microbial colonization, and in people living with diabetes. We used the library to link microbial *N*-acyl lipids, including histamine and polyamine conjugates, to HIV status and cognitive impairment. This resource will enhance the annotation of these compounds in future studies to further the understanding of their roles in health and disease and highlight the value of large-scale untargeted metabolomics data for metabolite discovery.

## Introduction

*N*-acyl lipids are signaling molecules consisting of two components: a fatty acid and an amine group, linked by an amide bond (**Figure 1A**). The previously described *N*-acyl lipids are involved in crucial biological functions, including immune homeostasis, building of fat mass levels, regulation of energy expenditure related to obesity, and they regulate other processes such as pain, memory, and insulin levels.^1–6^ Representative examples include *N*-oleoylethanolamine, which controls food intake, *N*-acyl taurine, which improves insulin sensitivity, and *N*-arachidonoyl 3-OH-γ-aminobutyric acid, which regulates calcium-dependent voltage channel function.^7,8^ Other *N*-acyl lipids, such as *N*-acetyl cysteine and N^α^-lauroyl-L-arginate, are used as an FDA-approved drug and a food ingredient, respectively. *N*-acetyl cysteine has antioxidant and anti-inflammatory properties and is used to block acetaminophen poisoning, as well as to break up mucus in respiratory diseases.^9^ On the other hand, N^α^-lauroyl-L-arginate acts as an antimicrobial agent, inhibiting bacteria, yeasts, and molds in food products.^8^ These are only a few examples of *N*-acyl lipids, but these molecules are chemically very diverse. LIPID MAPS, one of the most comprehensive lipid structural databases,^10^ currently catalogs close to 400 *N*-acyl lipids comprising 76 different headgroups derived from primary amines or amino acids (**Supplementary Figure 1A,B**).

**Figure 1.**
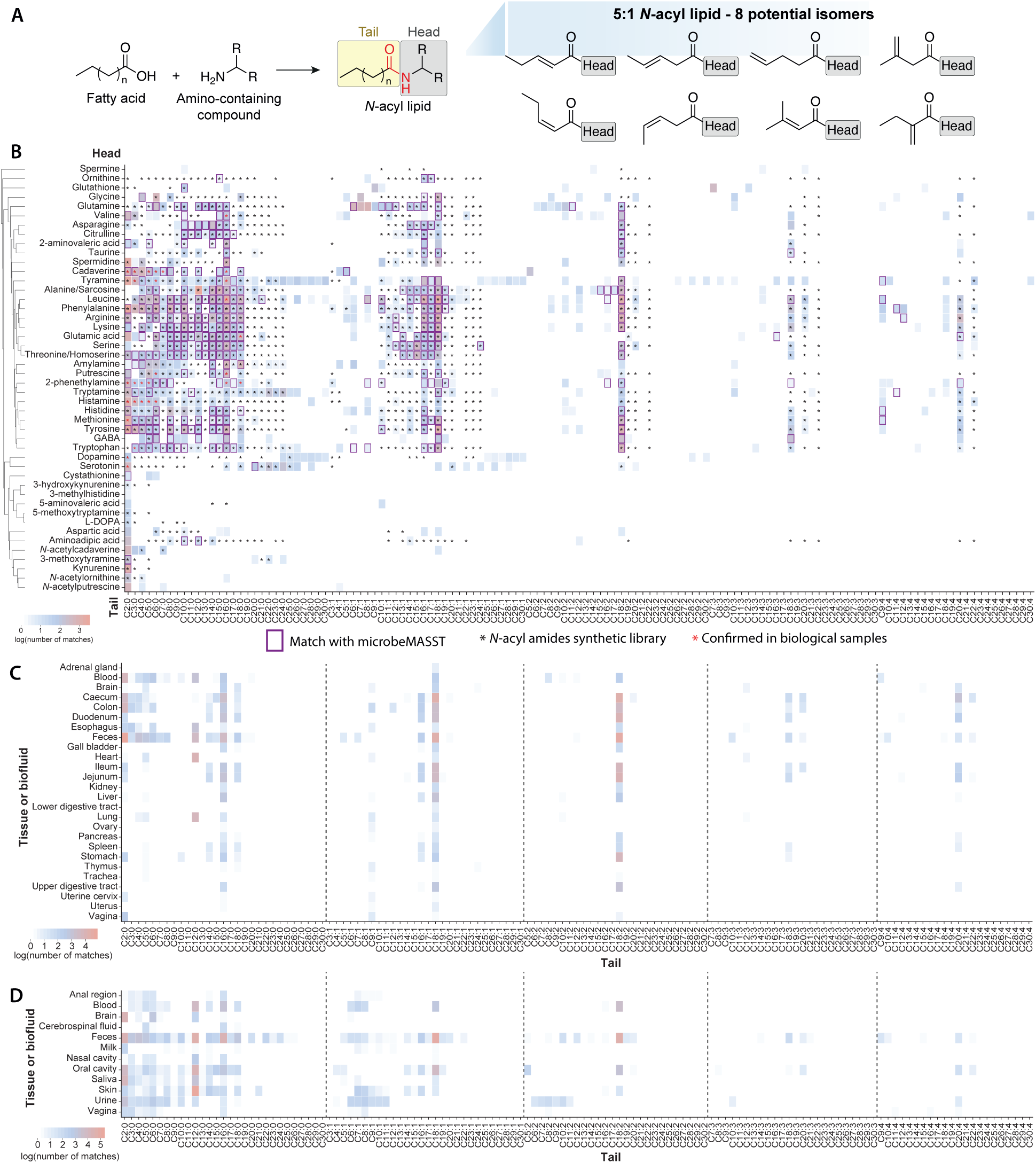
Repository-scale analysis of *N*-Acyl lipids in public mass spectrometry data and distribution among different tissues or biofluids. **(A)** *N*-acyl lipid definitions and isomers: this panel explains *N*-acyl lipids using a C5:1 tail example. A C5:1 lipid consists of a five-carbon fatty acid with one double bond. The image illustrates the possible isomers for this structure that can yield the same MS/MS spectrum. **(B)** Heatmap of *N*-acyl lipids: the heatmap shows 851 *N*-acyl lipids identified from public MS data in the MassIVE/GNPS repository using MassQL queries.^20^ Validation of the data was performed using cosine similarity (see **Supplementary Figure 1E**). Compounds found in microbial cultures are marked with purple squares, those matched with synthetic standards are indicated by black stars, and those confirmed by retention time with biological samples are shown with red stars. **(C)** and **(D)** Heatmaps showing distribution in tissues and biofluids: number of matches of different fatty acid chain lengths in tissues and biofluids with metadata available in ReDU^19^ for **(C)** rodent and **(D)** human-related public datasets. All heatmaps are shown as log values of the matches obtained from the repository, regardless of the headgroup. Icons were obtained from Bioicons.com.

Known *N*-acyl lipids can be identified through targeted mass spectrometry (MS) approaches,^11–14^ but both known and novel *N*-acyl lipids often go unreported in untargeted metabolomics data due to the lack of reference MS/MS spectra. We hypothesized that many *N*-acyl lipids relevant to biology exist within publicly available LC-MS/MS untargeted metabolomics data but remain unannotated due to the absence of relevant spectral libraries. Building on these efforts, we developed a novel strategy to create a reusable *N*-acyl lipid resource to reinterpret existing data from the untargeted metabolomics repository, GNPS/MassIVE. In this way, the biological function of *N*-acyl lipids in different contexts can be elucidated and we can ensure that future untargeted metabolomics studies will not overlook these important metabolites. Our approach leverages the reverse metabolomics strategy, where MS/MS spectra can serve as proxies for metabolites, which are then matched across public studies to contextualize their biological relevance.^14–22^

## Results

### Detection of *N*-acyl lipids in public data

To uncover the presence of *N*-acyl lipids and improve their detection in existing public untargeted metabolomics data, we developed the Mass Spec Query Language (MassQL) queries^20^ for 8,256 different *N*-acyl lipids with 64 amines and amino acids as headgroups (**Supplementary Table 1**, **Supplementary Figure 1C,D**). We applied these queries to filter MS/MS data from the GNPS-based untargeted metabolomics data (2,706 datasets as of January 2024, and includes ∼1.2 billion MS/MS spectra). The fragmentation-based queries were confined to 2- to 30- carbon fatty acids with up to four unsaturations. This range was selected because these fatty acids fragment predictably due to the limited presence of internal fragments, making it more straightforward to develop specific queries for which we could keep low false discovery rates (**Supplementary Methods, Supplementary Figure 1C,D**). Of the 64 headgroups we created queries for, 41 have not been documented in comprehensive curated lipid structure databases such as LIPID MAPS,^10^ LipidBANK,^23^ or SwissLipids,^24^ making their existence and/or prevalence in biology, including human biology, unclear (**Supplementary Figure 1A,B**).^25^

Of the 64 amines and amino acid headgroups, we found that 46 were linked with 2- to 30-carbon fatty acids in public data (**Figure 1B**). These represented 851 compounds of the theoretically 8,256 possible candidates from our initial MassQL searches (**Figure 1B, Supplementary Table S1**). A reusable MS/MS spectral library was created as a resource to enable other researchers to investigate *N*-acyl lipids in mass spectrometry-based metabolomics studies in the future. 552 spectra were confirmed to match their MS/MS using reference MS/MS of standards created using combinatorial organic synthesis^14,26^ (**Figure 1B**). These represent level 2 or 3 annotations according to both the 2007 Metabolomics Standards Initiative and the 2014 Schymanski rules for untargeted metabolomics annotation.^27,28^ In the absence of physical samples, this is the highest level of annotation currently possible for annotating MS/MS data in public data.

The most frequently detected lipid conjugate was acetylation (C2).^25^ While saturated carbons were the most common, an unexpected finding was the prevalence of both saturated and unsaturated C3-C6 short-chain fatty acid-derived *N*-acyl lipids, which are rarely reported in the lipid structural databases (**Supplementary Figure 1B**).^10,24,25^ Irrespective of fatty acid length, the saturated fatty acid containing *N*-acyl lipids were detected most frequently - followed by one, two, three, and four unsaturations. The most common fatty acids linked to *N*-acyl lipids were C18:1 and C16:1 for one double bond, and C18:2, C18:3, and C20:4 for two, three, and four unsaturations, respectively. Very long chain-linked *N*-acyl lipids are less frequently observed. Even-carbon lipid chains accounted for 87% of matches (**Supplementary Table S1**). Tyramine had the highest number of different fatty acid attachments, followed by leucine, phenylalanine, and tryptamine. Glutamine was associated with rare C8-C18 lipids, while tyramine, tryptamine, dopamine, and serotonin had rare C20-C30 lipid attachments (**Figure 1B**).

With the *N*-acyl lipids MS/MS spectra obtained using MassQL,^20^ we performed a MASST^18,29^ search against the entire GNPS repository to link the retrieved spectra to their biological associations. We obtained 356,542 MS/MS spectra from 61,833 files across 950 datasets, highlighting the widespread detection of *N*-acyl lipids in untargeted metabolomics studies. As little is known about the biology associated with *N*-acyl lipids, we leveraged the reverse metabolomics strategy^14^ to understand their presence in rodents, and humans, and their distribution across organs, biofluids, and other sources such as food, plants, or microbial cultures. By considering additional metadata curated with controlled vocabularies using the ReDU^19^ infrastructure in GNPS^21^, we could categorize *N*-acyl lipids detected in tissues and biofluids from humans and rodents, representing 435 and 259 *N*-acyl lipids, respectively. The tissue and biofluid distribution in rodents and humans, including the number of MS/MS spectra, unique *N*-acyl lipids headgroups, and different acyl chain lengths are depicted in **Supplementary Figure 2A,B**. The most frequently observed chain lengths in both humans and rodents were C2, C12, C16, and C18, as illustrated in **Figures 1C** and **1D**. Odd-chain lipid chains were also detected in both human and rodent datasets with C3:0 (propionate) and C5:0 (valerate), both classified as short-chain fatty acid-derived molecules, being the most frequently detected among them. In rodents, C3:0 was primarily observed in the colon, caecum, esophagus, and feces, while C5:0 was mostly found in feces and blood. In humans, C3:0 was detected in saliva, the vagina, and feces, while C5:0 was present in the oral cavity, urine, blood, and cerebrospinal fluid, in addition to feces. The most common head groups identified in both humans and rodents were phenylalanine, spermidine, (iso)leucine, and alanine/sarcosine (**Supplementary Figure 2C,D**). The data suggests that *N*-acyl lipids occupy specific body niches. Aspartic acid, aminoadipic acid, and spermidine lipids were primarily found in the brain and rarely in other body locations. Spermidine-conjugated lipids appeared frequently in saliva, while glutamine *N*-acyl lipids were more common in blood, skin, and urine. Tyrosine-conjugated lipids, however, were almost exclusively detected in human milk.

Out of the 851 *N*-acyl lipids, 347 were detected in data from microbial cultures using microbeMASST^15^ (**Figure 1B**, **Figure 2A,B**, **Supplementary Table S1**). The most commonly observed *N*-acyl lipids in these microbial monocultures had phenylalanine, leucine, and tyrosine as headgroups (**Figure 2A**), with an overall predominance of even-chain lengths (**Figure 2B**). Additionally, 167 and 243 of the 851 candidate *N*-acyl lipids were detected in plant and food datasets, respectively (**Figure 2C, Supplementary Table S2**). This distribution stratified by lipid chain length revealed that short, medium, and long-chain conjugates are predominantly detected in human, microbial, and rodent-related datasets, while very long-chain *N*-acyl lipids are observed almost exclusively in plants and foods (**Figure 2D**, **Supplementary Figure 3A-D**). These differences in *N*-acyl lipids found in food and plant data compared to microbial cultures, rodent, and human datasets suggest they may be consumed through diet but also produced by the microbiota.

**Figure 2.**
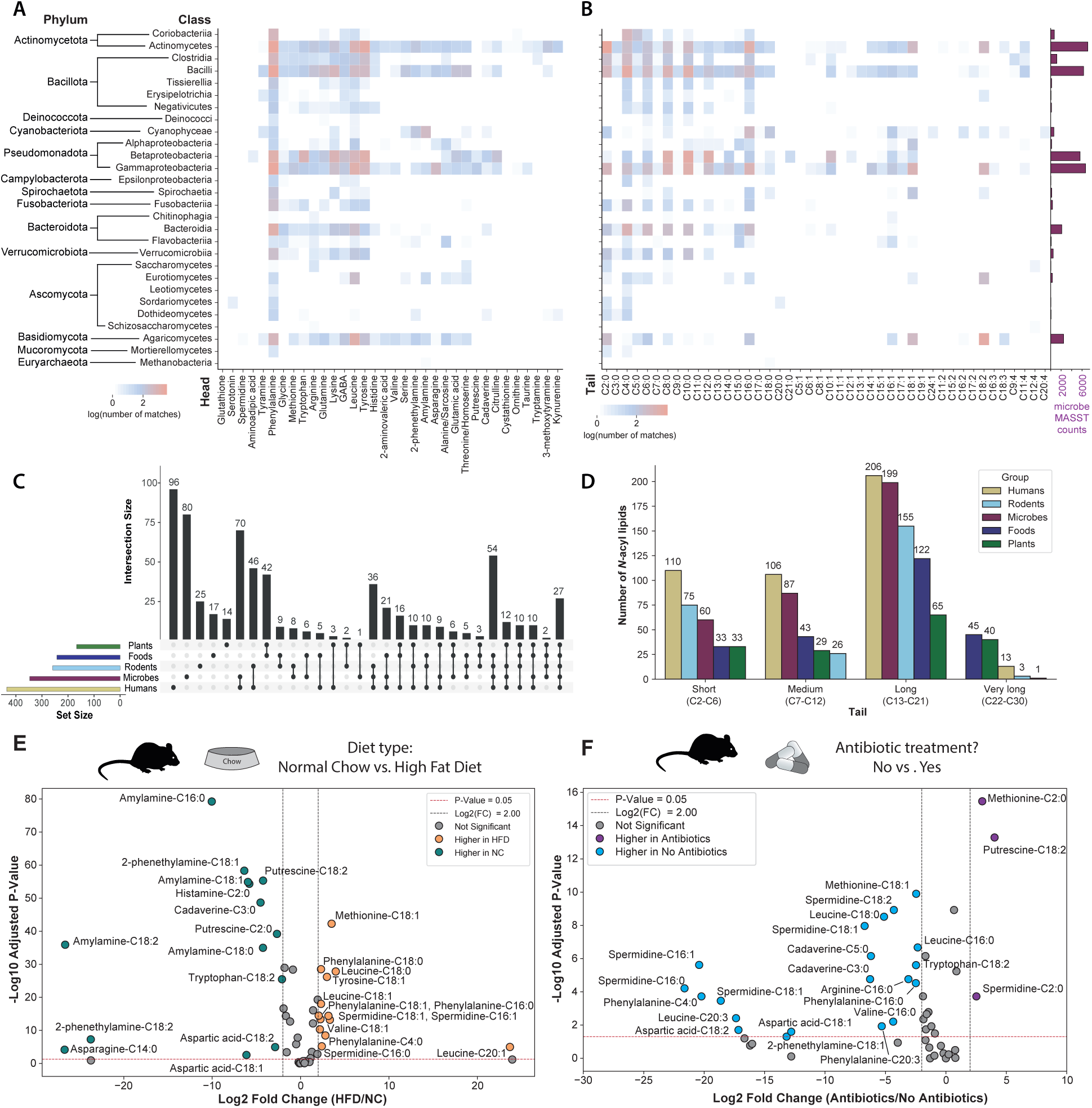
Evidence of microbial origins of *N*-acyl lipids. Heatmaps depict the distribution of different headgroups (**A**) and tails (**B**) across various microbial classes, with barplots showing the total counts for each class in microbeMASST.^15^ The Y-axis was taxonomically ordered according to the NCBI Taxonomy ID, while the X-axis was clustered using the Braycurtis metric for the headgroups, or in ascending order (in number of carbons and unsaturations) for the tails. **C)** UpSet plot of *N*-acyl lipid distribution: This plot highlights the distribution of *N*-acyl lipids across different datasets, including human-related, rodent-related, microbial monocultures, plant-, and food-associated data. **D)** Distribution of *N*-acyl lipid chain lengths: This summary shows the prevalence of short, medium, long, and very long chain *N*-acyl lipids in public data. Note that the exact location and cis/trans configurations of double bonds cannot be determined from the current queries, which are annotated at the molecular family level according to the Metabolomics Standards Initiative.^27^ **E** and **F)** Volcano plots of mouse fecal pellets from a dataset publicly available (GNPS/MassIVE: <u>MSV000080918</u>)^30^ showing *N*-acyl lipids up-regulated and down-regulated upon different diets (**E**) and antibiotic treatment (**F**). The significant thresholds are marked by dotted lines in the volcano plot (p < 0.05 and log2(FC) > 2 or <2). Differential compounds between the groups were evaluated using the non-parametric two-sided Mann-Whitney U test, and p-values were corrected for multiple comparisons using the Benjamini-Hochberg correction. Icons were obtained from Bioicons.com.

This hypothesis was further evaluated by the analysis of a public dataset of small intestine and colon samples, where germ-free (GF) mice were colonized with a conventional gut microbiota (Specific Pathogen Free, SPF), or monocolonized with Segmented Filamentous Bacteria (SFB), or other gut commensal strains.^31,32^ In addition, we conducted another culturing experiment with human-derived microbiota to enable MS/MS and retention time matching. Both datasets revealed a mixture of both consumption and production of *N*-acyl lipids, providing additional evidence that the microbiota regulates *N*-acyl lipid levels. These results are alsoconsistent with a recent report of a *Faecalibacterium prausnitzii* hydrolase that has both amide hydrolase and *N*-acylation function.^33^

The *N*-acyl lipid profile in the small intestine and colon differed in mice colonized with conventional microbiota (SPF) or monocolonized with SFB compared to GF mice. In addition, the reanalysis of other monocolonized mouse samples revealed that short-chain fatty acids generally increased, while those conjugated with longer chains decreased compared to GF mice, with cases of microbe-specificity, supporting the hypothesis that microbes may be involved in *N*-acyl lipids production (**Supplementary Figure 3E**). Culturing 71 commensal bacteria from the human gut also revealed their ability to make *N*-acyl lipids and provided additional support for this hypothesis.^34^ Since the vast majority of these microbes are not yet part of microbeMASST, this approach provided both orthogonal evidence and experimental validation of microbial-linked *N*-acyl lipids. We obtained 50 MS/MS matches to the *N*-acyl lipids resource, with 38 corresponding to *N*-acyl lipids conjugated to short-chain fatty acids (**Supplementary Figure 3F, Supplementary Figure 4A,B**). We observed that short-chain *N*-acyl lipids increased compared to the culture media, while longer chains (C8-C12) generally decreased, except for ornithine-C17:1, and leucine and methionine-C9:4, suggesting that the microbiota is able to produce many of these *N*-acyl lipids conjugated to short-chain fatty acids.

To assess the presence of *N*-acyl lipids and their potential changes under different biological conditions, we performed in-depth analyses using our newly created library on public datasets that had expanded metadata. Reanalysis of datasets on diabetes (type I), various stages of forensic human body decompositions,^35^ and diet and effect of antibiotics in colorectal cancer^30^ revealed the presence of many *N*-acyl lipids based on matching their MS/MS against the MS/MS *N*-acyl lipids resource (**Figure 2E,F**, **Supplementary Figure 3G-M**). Peak intensity analysis against the available metadata revealed that shorter-chain *N*-acyl lipids were decreased in the diabetic group (urine from humans) (**Supplementary Figure 3G**), longer-chain fatty acids *N*-acyl lipids increased upon cadaver decomposition (skin swabs from humans and soil) (**Supplementary Figure 3H-K, Supplementary Figure 4D,E**), and overall *N*-acyl lipids levels were altered by diet (feces of mice on normal chow vs. high fat diet) (**Figure 2E**). Mice on a normal chow had a higher abundance and variety of short-chain fatty acid-derived *N*-acyl lipids compared to mice on a high-fat diet. Conversely, mice on a high-fat diet showed increases in *N*-acyl lipids conjugated to longer-chain fatty acids (**Figure 2E**, **Supplementary Figure 3L**). Intriguingly, most of those same longer-chain fatty acid conjugates that are observed in the high-fat diet are no longer detected upon treatment with an antibiotic cocktail (**Figure 2F, Supplementary Figure 3M**), providing additional evidence linking the production of many of the *N*-acyl lipids conjugates to the microbiome and diet. After generating and validating the *N*-acyl lipid resource with published datasets and with the new knowledge that many *N*-acyl lipids are made by the microbiota, we next set out to demonstrate its utility in a new human research study.

### Demonstrating the utility of the *N*-acyl lipid resource - *N*-acyl lipids in relation to HIV, immune and cognition status

To further demonstrate the utility of our newly created *N*-acyl lipid MS/MS library, and to provide a case study on how to leverage this resource, we used it to annotate *N*-acyl lipids in an ongoing study in our laboratory aimed at understanding the effect of the microbiome on cognition in people infected with the human immunodeficiency virus (HIV). This cohort included stool data collected from both people with HIV (PWH) and people without HIV (PWoH) who had also completed neurocognitive evaluations as part of NIH-funded studies conducted at the UC San Diego HIV Neurohevavioral Research Program, primarily the HIV Neurobehavioral Research Center (HNRC). More than 50 matches to MS/MS spectra of *N*-acyl lipids were obtained, and we observed higher levels of histamine *N*-acyl lipids, particularly those conjugated with short-chain fatty acids, in PWH compared to PWoH (**Supplementary Figure 5A**). In pairwise comparisons of specific *N*-acyl lipids, histamine-C2:0, histamine-C3:0, and histamine-C6:0 were higher in PWH (Mann-Whitney U test, p-values of 0.003, 0.003, and 0.042, respectively). Besides that, histamine-C4:0 and histamine-C5:0 also showed a higher trend in PWH. All other *N*-acyl histamines, including those not initially searched for with the MassQL query but identified through molecular networking,^36–38^ were found in higher average levels in samples of PWH compared to PWoH (**Supplementary Figure 5B**). However, none of these lipids reached significance at the selected statistical threshold of p < 0.05.

Histamine conjugates were linked to HIV status, while polyamine *N*-acyl lipids were associated with neurocognitive impairment status (impaired vs. unimpaired). Specifically, cadaverine and putrescine *N*-acyl lipids, particularly those with short acyl chains, were elevated in the impaired group compared to the unimpaired group (Mann-Whitney U test, p-values from 0.001 to 0.04, **Supplementary Figure 5C,D**). Further, analyses using a linear mixed-effects model, with fixed covariates of HIV status and neurocognitive impairment status, while treating the subject as a random effect, suggest that histamine-C2:0 and histamine-C3:0 continue to be positively associated with HIV status and acylated polyamines were associated with neurocognitive impairment (**Figure 3A**). We also observed a trend where the histamine conjugates with C2, C3, C4, and C5, were negatively associated with CD4^+^/CD8^+^ T cell ratio in PWH, which is an indicator of immune system homeostasis.^39^ In contrast, polyamines, particularly cadaverines linked to C2, C3, C5, C6, and C7, tended to show a positive correlation with the CD4^+^/CD8^+^ ratio in PWH (**Figure 3B**). Additionally, we explored the relationships between *N*-acyl lipids and plasma HIV RNA viral loads in the PWH. We found that *N*-acyl lipids with short acyl chains were positively associated with higher viral loads, while those with longer acyl chains were linked to lower viral loads (**Supplementary Figure 5E**). To validate their identities, we matched retention time and MS/MS in comparison to pure synthetic standards for 13 of these short-chain fatty acid-derived *N*-acyl lipids that are associated with HIV status (histamine-C2:0, C3:0, C4:0, C5:0, and C6:0), neurocognitive impairment status (cadaverine-C2:0, C3:0, C5:0, C6:0, and C7:0), and included dopamine-C2:0, serotonin-C2:0, and tryptophan-C3:0 in this validation of the annotations neurotransmitters derivatives - even though they did not associate with neurocognitive impairment (**Figure 3C**). All of the compounds matched both the retention times and the MS/MS spectra in the fecal samples, confirming their presence in the samples (**Supplementary Figure 5F,G**). Quantification revealed that many of these can be present in high µM concentrations (**Supplementary Table S3**). Although we do not yet understand the biology behind this variability, and many samples had concentrations below the limit of quantification, the highest concentrations of *N*-acyl histamines that we quantified were 93.8, 20.7, 7.0, and 2.7 ng/g of fecal sample for histamine-C2:0 through histamine-C5:0, respectively. Additionally, for the *N*-acyl cadaverines, we found the concentrations to be as high as 350.4, 126.7, 36.7, and 1.3 ng/g for cadaverine-C2:0, cadaverine-C3:0, cadaverine-C5:0, and cadaverine-C6:0, respectively. Dopamine-C2:0 was also quantified, with levels ranging from 0.0008 to 3.2 ng/g. While histamine-C6:0, cadaverine-C7:0, serotonin-C2:0, and tryptophan-C3:0 were matched with retention times and MS/MS, their concentrations were too low to be accurately quantified in the samples.

**Figure 3.**
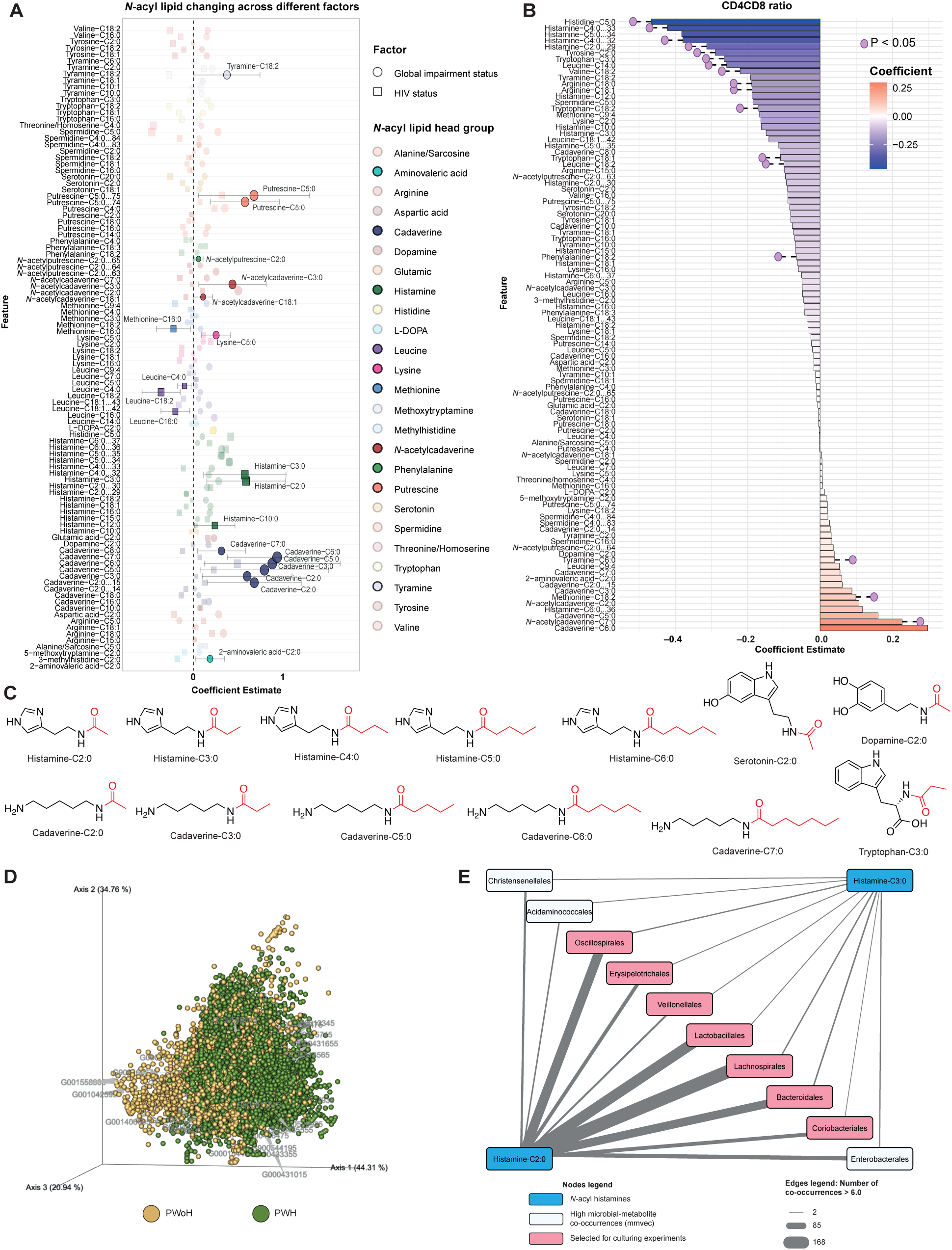
*N*-acyl lipids are correlated with HIV and neurocognitive impairment status. (**A**) Forest plot illustrating the coefficient estimate of a linear mixed-effects model for individual *N*-acyl lipid species, with fixed covariates of HIV status (PWH, *n* = 226; PWoH, *n* = 87) and neurocognitive impairment status (impaired, *n* = 151; unimpaired, *n* = 162), accounting for random effects within individual samples/visit. Filled circles (HIV status) and squares (neurocognitive impairment status) with corresponding confidence intervals represent significant *N*-acyl lipid species. Faded circles and squares depict non-significant species. Each color represents a different headgroup. (**B**) Bar plot showing the correlation coefficients of association between CD4/CD8 ratio and various *N*-acyl lipids in a subset of the PWH (*n* = 171) with available metadata. Red bars represent positive correlations, while blue bars represent negative correlations, as determined by linear regression models. The p-values shown are nominal; adjusted p-values (corrected for multiple comparisons using the Benjamini-Hochberg method) are available in **Supplementary Table S3**. (**C**) Structures of all *N*-acyl lipids confirmed in this study with pure synthetic standards. (**D**) Microbe-metabolite co-occurrence biplot obtained from mmvec^40^ analysis of the HNRC sample. Spheres represent ions of molecules, while arrows represent microbes. Spheres were colored based on which group (PWH vs. PWoH) each ion feature was most abundant in. Small angles between the arrows indicate microbes co-occurring with each other, and spheres close in the plot represent features co-occurring. Arrows pointing toward a group of molecules indicate microbe-molecule co-occurrence. This biplot shows the 30 most important OTUs (higher vector magnitude). (**E**) Network of the microbial taxonomic orders with co-occurrences > 6.0 and shared between histamine-C2:0 and histamine-C3:0. Nodes colored in pink are the orders selected for culturing experiments.

### Microbial producers of HIV-associated histamine *N*-acyl lipids

Samples from the HNRC also underwent metagenomic sequencing, allowing us to perform correlation analyses to identify microbes potentially responsible for producing the histamine and polyamine *N*-acyl lipids associated with HIV and neurocognitive status. Previous microbial cultures from this study produced cadaverine-C2:0 and cadaverine-C4:0 (**Supplementary Figure 3F**). MicrobeMASST searches also confirmed that cadaverine *N*-acyl lipids have been observed in microbial monocultures (**Figure 1B**). However, no microbial *N*-acyl histamines were detected in either public data or our experiments, raising the question of whether histamine conjugates are microbially produced, and if so, which microorganisms may be responsible for their production.

To investigate this further and identify microbes potentially associated with N-acyl histamines, we conducted a multiomic microbe-metabolite co-occurrence analysis using mmvec.^40^ We observed a strong trend of distinct microbe-metabolite co-occurrences between PWH and PWoH (**Figure 3D**). Higher microbial-metabolite co-occurrence probabilities were observed for histamine-C2:0 and histamine-C3:0 (**Supplementary Table S3**). Ten microbial taxonomic orders also stood out for presenting several organisms that resulted in high co-occurrence probabilities with both histamine-C2:0 and histamine-C3:0, with histamine-C2:0 exhibiting more high co-occurrences than histamine-C3:0 (**Figure 3E**).

Based on the multiomics analysis and availability of strains, we selected nine bacterial strains from these microbial orders for culturing and supplemented the media with histamine, cadaverine, and putrescine. After 72 h of culturing, we analyzed the samples using LC-MS/MS and matched them against the *N*-acyl lipids library (**Figure 4**). We identified histamine-C2:0, histamine-C3:0, cadaverine-C2:0, and cadaverine-C3:0 in the cultures at 72 h, whereas these compounds were not detected at 0 h post-addition. This finding confirmed that some microorganisms were capable of producing these *N*-acyl lipids. Specifically, cadaverine-C3:0 was observed in cultures of *Collinsella aerofaciens* ATCC 25986 and *Prevotella buccae* D17, while cadaverine-C2:0 was detected in extracts from these two microbes as well as in *Catenibacterium mitsuokai* DSM 15897 and *Holdemanella biformis* DSM 3989. *Catenibacterium mitsuokai* DSM 15897 produced only histamine-C2:0, whereas *Collinsella aerofaciens* ATCC 25986, *Holdemanella biformis* DSM 3989, and *Prevotella buccae* D17 produced both histamine-C2:0 and histamine-C3:0. *Dorea longicatena* DSM 13814 produced only histamine-C3:0. *Collinsella aerofaciens* ATCC 25986 produced the highest amount of histamine-C2:0, with a concentration of 1.905 ± 0.302 µM in the extracts (**Figure 4A, Supplementary Table S4**). The highest levels of histamine-C3:0 were observed in *Prevotella buccae* D17, with a concentration of 0.358 ± 0.016 µM (**Figure 4B, Supplementary Table S4**). Cadaverine-C2:0 and cadaverine-C3:0 were confirmed to be produced by specific microbes by MS/MS and retention time matching, but these compounds were present in lower concentrations in the extracts and could not be accurately quantified.

**Figure 4.**
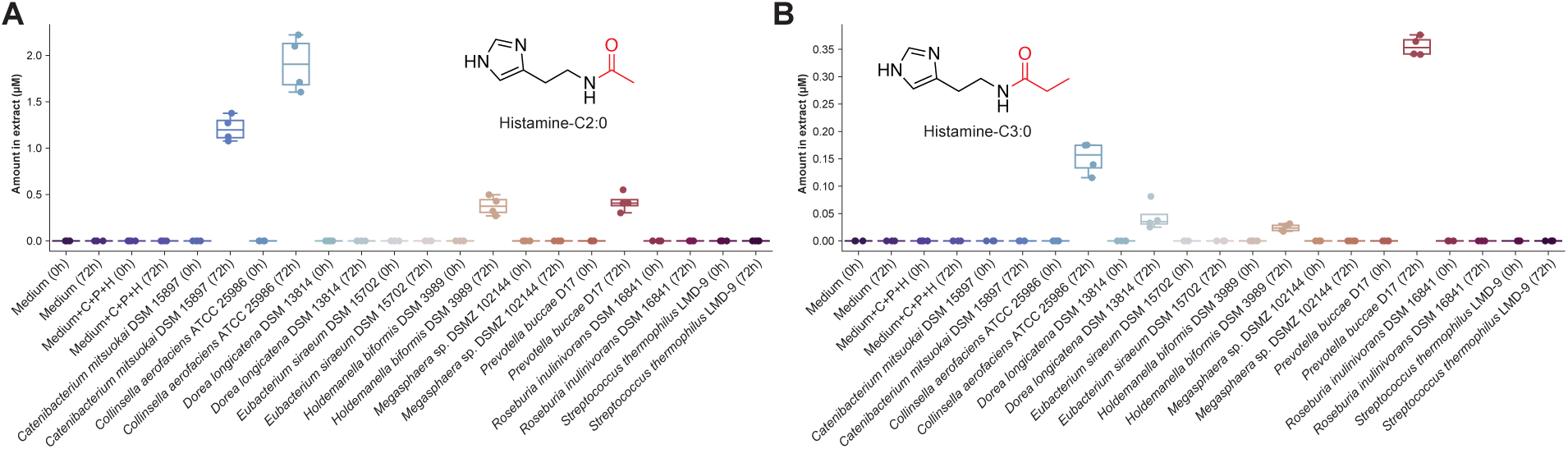
Evidence of microbial production of *N*-acylated histamines. Concentrations of **A)** histamine-C2:0 and **B)** histamine-C3:0 in microbial extracts. Values in the y-axis represent the amount of these compounds in micromolar (µM) concentrations in the extracts. Cadaverine (C), putrescine (P) and histamine (H) were added to the medium.

## Discussion

Despite their infrequent description or annotation in metabolomics data and literature, *N*-acyl lipids are quite prevalent, as revealed by our reverse metabolomics analysis using the MassQL-generated MS/MS reference resource. A significant portion of the *N*-acyl lipids identified in this study were derived from short-chain fatty acids. Free short-chain fatty acids are a key and extensively studied class of molecules produced at the microbiota-diet interface.^41^ While primarily produced in the gut, these fatty acids can impact distant organs such as the liver, lungs, urogenital tract, and brain.^42–44^ They play a role in immune regulation, affecting T-cell functions, CD4^+^/CD8^+^ levels, and are implicated in health conditions like disrupted intestinal barrier function, and diseases, including autoimmune disorders, diabetes, and HIV.^41,45,46^

Given the short-chain fatty acids prominence in microbiome research, it was surprising to find such a large panel of *N*-acyl lipids, many derived from short-chain fatty acids, that had not been previously documented in lipid structural data resources. Our study demonstrated that these *N*-acyl lipids are detected in data from sites distant from the gut. Their levels are influenced by factors such as dietary changes, antibiotic use, and health conditions affecting the microbiome, such as diabetes, and other microbiome-mediated processes such as decomposition. Additionally, analysis of publicly available data, along with microbial culturing experiments conducted in this study, showed that individual cultures can produce, in a microbe-specific manner, certain *N*-acyl lipids when both the amine headgroup and lipid substrates are present.

Data from fecal samples of people with HIV revealed a substantial number of microbially produced short-chain fatty acid-derived *N*-acyl lipids—an observation not previously identified despite numerous metabolomics studies on PWH.^47–54^ This discovery was made possible by the N-acyl lipid MS/MS resource created in this work. We found that microbially produced short-chain fatty acids linked to polyamines and histamine were associated with plasma HIV RNA viral load and CD4^+^/CD8^+^ levels in PWH. This study uncovered several *N*-acylated lipids related to HIV status, including histamine conjugates, while many polyamine-derived *N*-acyl lipids were associated with neurocognitive impairment status in both people with and without HIV in this cohort.

Although limited information is available on histamine-containing *N*-acyl lipids, it is known that the non-acylated histamine itself, produced by macrophages, is increased in people with HIV.^55^ Thus, one can hypothesize that the production of the *N*-acylated histamines may require not only the availability of histamine but also the short-chain fatty acids and the right organisms. Indeed, organisms such as *Prevotella*, which are commonly enriched in PWH,^56^ can produce propionate from succinate^57^ and have the ability to couple this to histamine. Beyond HIV populations, very little is known about the short-chain fatty acid-histamine conjugates. The C2 and C3 *N*-acyl histamines were previously found to be elevated in the urine of patients with intestinal disorders, the histamine-C6 was found to be very modestly cytotoxic while related molecules that have longer chain fatty acids conjugated to them act on peroxisome proliferator-activated receptor-α (PPAR-α).^58–60^ PPAR-α protects from HIV-related systemic inflammation and improves intestinal barrier function.^61,62^ We did not find biological reports for the C4-C5 histamine conjugates, and it is not yet known if these specific *N*-acyl lipids also provide such protective effects.

There is a strong connection between HIV disease and polyamines. Polyamines, such as cadaverine, protect the HIV virion and sperm from the acidity in the vaginal tract and increase infectivity.^63^ Polyamines are detected in higher quantities and affect T_reg_ cell dysfunction in people with HIV.^48,64,65^ Intriguingly, polyamine metabolism plays a crucial role in maintaining the integrity of helper T cell lineage, which is crucial in regulating inflammation and maintaining immune tolerance.^66,67^ As with the histamine conjugates, there is also limited information on polyamine conjugates and HIV disease or other health conditions. This includes the cadaverine *N*-acyl lipids, except for the commonly measured C2-conjugate, which has been associated with cancer and other health conditions, such as in the urine of individuals with Alzheimer’s disease and has been shown to reduce the aggressiveness of breast cancer in rodents.^68–70^ The cadaverine-C3, also known as *N*-propionyl cadaverine, has been shown to reach the brain of rats *in vivo* but also, as shown *in vitro*, depresses electrically stimulated dopamine release from the neostriatum from rats at concentrations in the nM range.^71–73^ We did not find biological reports for the C4-C6 cadaverine, despite recent studies highlighting the discovery of microbiome-derived polyamines.^22,74–76^ We found polyamine *N*-acyl lipids, especially cadaverine short-chain fatty acid conjugates, are associated with impairment status in this study, a new finding, although it is established that other polyamines, acetyl-spermidine, and unconjugated putrescine, are biomarkers for HIV-associated neurocognitive disorders,^77^ this is not known for the *N*-acyl-cadaverines.

Non-dietary histamine and cadaverine levels are reported to be inversely linked, and we see a similar trend for the *N*-acyl cadaverine and histamines and in relation to CD4^+^/CD8^+^ ratio. Although it is not yet known if *N*-acyl conjugates exhibit similar activities, cadaverine can potentiate histamine levels, possibly via competitive inhibition of histamine-degrading enzymes.^78–80^ This inverse relationship and the role of the *Prevotella-*derived production of short-chain fatty acid-linked histamine and polyamines and their role in HIV disease and HIV-associated neurocognitive impairment warrants more research. However, without the *N*-acyl lipid reference resource provided by this work, the observation that these microbial-derived molecules and their associations with HIV disease and HIV neurocognitive impairment would have remained hidden.

The unexpected discovery of hundreds of short-chain fatty acid-derived *N*-acyl lipids, not reported in structural lipid databases, highlights their widespread presence across all biofluids and organs for which data is available, despite most being produced by the microbiota. The identification of various structural family members opens an additional chapter in understanding the mechanistic and functional roles of short-chain fatty acids. This finding may even prompt a reinterpretation of how microbially produced short-chain fatty acids influence the production of *N*-acyl lipids, and consequently, a wide range of conditions, as they are only formed when both substrates are present and the appropriate microbe(s) are present to create the link. This resource has enabled the generation of numerous hypotheses regarding the functions of these *N*-acyl lipids, and we anticipate that fully elucidating their roles will require extensive research across many laboratories and thousands of studies.

While we provide signatures for 851 metabolites here, this is only the beginning. Many other amines are not covered in this study and they also may be linked to different fatty acids. Lipids containing other atoms, such as oxygen, nitrogen, or halogens, were not included and would require dedicated MassQL queries or the development of alternative detection strategies. Alcohols might also undergo similar structural diversification. Moreover, the diversity of lipids available for acylation extends well beyond the C2:0 to C30:4 range of lipids we examined. We anticipate that this resource will spur the development of additional ways to find *N*-acyl lipids and will help uncover additional biological and health associations. This may enhance our understanding of microbiome-mediated effects and potentially serve as easy-to-detect microbial biomarkers in precision medicine, given their prevalence. Finally, this resource captures the intersection of nutrient availability with microbial and host metabolism, warranting further exploration as regulators of the immune system.

### Limitations of the study

Users of this resource should consider three main limitations when making biological discoveries. First, while we have consistently matched the MS/MS of synthetic standards to MassQL recovered spectra, there have been instances where the match was to a different isomer. For example, in the cohort of the body decomposition study, there were compounds, such as the amylamine conjugates, that the MS/MS spectra matched the standard, but the retention times did not align, suggesting the presence of a different isomer instead, such as a branched chain in the acyl portion. Other headgroups can also have more than one position for the acyl attachment, which will also result in very similar MS/MS spectra. For the HIV study, all the pure *N*-acyl lipid standards matched the compounds present in the samples. Even though there were three nitrogen atoms available for the acyl substitution, the substitution of the acyl chain was observed in the primary amine group in all cases. Therefore, at the repository level search, it is advisable to refer to the number of carbons and double bonds in the lipid chain rather than the exact structure, as multiple isomers can correspond to the same family of molecules (see **Figure 1a**).

Secondly, our initial query was designed to capture the protonated ion forms of the molecules. However, many different ion forms, such as adducts, multimers, and in/post-source fragments, are commonly detected for any given molecule. The fragmentation patterns of other ion forms may differ and would require separate MassQL queries. A limitation of using other ion forms for query development is the scarcity of reference spectra to understand their fragmentation behavior. Nevertheless, once an annotation is made, it is possible to retrieve associated MS/MS spectra for different ion forms through peak shape and retention time alignments.^81^ Currently, this type of analysis is feasible only within a single dataset and not across all public data simultaneously.

Finally, it is crucial to note that our observations are based on the *N*-acyl lipid spectra detected in public-domain data. Biological associations can only be established when there is well-curated public (meta)data. Variations in underlying biological conditions—such as feeding time, health, circadian rhythm, and diet type—may affect concentrations and detectability in untargeted metabolomics. Moreover, mass spectrometry-based metabolomics data are highly sensitive to data acquisition parameters (e.g., mass spectrometer type, ionization technique, collision energies, chromatographic gradient) and sample preparation methodologies (e.g., storage conditions, extraction methods). Therefore, while the observed patterns and trends in *N*-acyl lipid distribution across various tissues and biofluids provide valuable insights, they should be interpreted with these considerations in mind and use the results to formulate testable hypotheses.

## Acknowledgments

We thank the support by NIH (NIDDK) for the development of tools for structure elucidation R01DK136117, the Collaborative Microbial Metabolite Center U24DK133658, and the HIV Neurobehavioral Research Center (HNRC) is supported by Center award P30MH062512 from NIMH. This work was further supported by BBSRC-NSF award 2152526. Research reported in this publication was supported in part by the National Center for Complementary and Integrative Health of the NIH under award number F32AT011475 to N.E.A., the Maternal and Pediatric Precision in Therapeutics project P50HD106463. X.S. was supported by the National Key R&D Program of China 2022YFA0807300, 2023YFA1800200, NSF of China 32270945, STCSM 22ZR1468700, and 22140902400. S.L. was supported by Research Council of Finland funding (no. 363417 to S.L.). J.J.K. was supported by the NIH CTSA grant UL1TR001442 and the UCSD Microbiome Seed Grant. M.R. was supported by NIH grant R37AI126277. G.T.W. was supported by NIH training grant T32AI007036. G.J.N. was supported by NIH fellowship F31AI186410. We also thank Dr. Jessica L. Metcalf for the supervision of the human cadaver decomposition study and Dr. Robert Heaton for the participation in the development of the clinical cohort of human immunodeficiency virus (HIV) infection.

## Author contributions

P.C.D. conceptualized the project. H.M-R. and M.F. developed the MassQL queries and performed the repository-scale searches. H.M-R. created the N-acyl lipids library. H.M-R, S.L., W.D.G.N., L.K., P.R., H.N.Z, and P.W.P.G. performed data analysis. V.C-L., G.W., G.N., L.C., C.W., and S.R. performed microbial incubation experiments. K.Z., M.R., and L.B. supervised the culturing experiments. H.M-R., V.C-L., W.D.G.N., J.Z., and M.J.M. acquired LC-MS/MS data. V.D., A.P., K.V., I.M., C.L., A.S., M.S., Y.L., and D.P. performed the combinatorial synthesis reactions. P.C.D. and D.S. supervised the synthesis. M.S.A., R.J.E., D.J.M, J.E.I., D.F.Jr, and S.L. developed the clinical cohort of human immunodeficiency virus (HIV) infection. M.S.A. and C.X.W. assisted with data interpretation. Y.E.A. performed database searches. M.W. provided support for MassQL searches. X.S. and D.K. supervised the monocolonized mice study. J.K. and S.K. supervised the diabetes study. Z.B. supervised the human decomposition study. R.K. supervised sample handling and DNA data acquisition for the HNRC cohort. H.M-R. and P.C.D. drafted the manuscript. P.C.D. acquired funding and supervised this project. All authors reviewed and edited the manuscript.

## Supplementary figure titles and legends

**Supplementary Figure 1. Distribution of *N*-acyl lipids in structural databases and mass spectrometry repository searches, related to Figure 1. A)** Diversity and relative frequency of *N*-acyl lipids headgroups and **(B)** lipid chain lengths documented in LIPID MAPS. This analysis excludes ceramide acylations. **C)** *N*-acyl lipid query strategy: representative MS/MS spectrum of phenylalanine-C10:0 (CCMSLIB00011435104) and phenylalanine-C16:0 (CCMSLIB00011435452). The spectra show nearly identical fragmentation patterns enabling the creation of the MassQL query to retrieve the MS/MS spectra of this family of lipids. **D)** MassQL query for phenylalanine headgroup where we initiate to return all MS/MS spectra (in yellow) that fulfill the following criteria: the precursor ion has to match one of the expected precursor *m/z* values specified (gray), as well as the most diagnostic *m/z* fragments of the head portion (blue and pink) with their indicated error tolerances and minimum relative intensities. **E)** Strategy followed to create the *N*-acyl lipids library and expand to biological interpretations. (I) MassQL queries were designed and run against the Orbitrap datasets in the GNPS/MassIVE repository. (II) The spectra were clustered using MSCluster to reduce redundancy. (III) A cosine similarity filter was applied to keep the higher confidence *N*-acyl lipids spectra. (IV) The clustered spectra were searched using FASST searches against the whole repository (including Orbitrap and QToF datasets), and human and rodent-related datasets were tagged using ReDU, and microbial, plant, and food-related datasets were also tagged using domain-specific MASSTs. (V) The spectra retrieved from the FASST searches were filtered to keep the matches in which the raw (unfiltered) spectra resulted in cosine similarity above 0.7. (VI) Summary of the results obtained with this workflow. Icons were obtained from Bioicons.com.

**Supplementary Figure 2. Distribution of *N*-acyl lipids obtained from FASST searches among different tissues or biofluids, related to Figure 1**. Summary of the occurrences in the public domain in (**A**) human and (**B**) rodent-related datasets. Heatmaps show the distribution of the number of matches grouped by headgroup in different tissues and biofluids with metadata available in ReDU for (**C**) human and (**D**) rodent-related public datasets. All heatmaps are shown as log values of the matches obtained from the repository. Icons were obtained from Bioicons.com.

**Supplementary Figure 3. *N*-acyl lipids chain length diversity, evidence of microbial *N*-acyl lipids, and reanalysis of public datasets, related to Figure 2**. Distribution of *N*-acyl lipids in public data stratified by chain length classes. Upset plots show the number of unique *N*-acyl lipids attached to (**A**) short, (**B**) medium, (**C**) long, and (**D**) very long chain fatty acids. (**E**) Reanalysis of a public dataset of monocolonized GF mice (GNPS/MassIVE: <u>MSV000088040</u>, deposited in 2021)^31,32^. Heatmap log2 fold changes (FCs) of the *N*-acyl lipids matches in colon and small intestine samples of monocolonized mice relative to germ-free (GF) mice. Values of the diet, Specific Pathogen Free (SPF) mice, and of mice colonized with Segmented Filamentous Bacteria (SFB) are also shown. Red cells indicate compounds that are increasing relative to GF, while blue cells indicate compounds that are decreasing relative to GF mice. The x-axis is taxonomically ordered according to the NCBI Taxonomy ID. (**F**) Heatmap showing the log2 fold change of N-acyl lipids matches in microbial monocultures of gut commensal microbes relative to the culture media. Red cells indicate compounds that are increasing, while blue cells indicate compounds that are decreasing relative to the media. The x-axis is taxonomically ordered according to the NCBI Taxonomy ID. (**G**) Peak area abundances of *N*-acyl lipids annotated in a public dataset (GNPS/MassIVE: <u>MSV000082261</u>) from urine samples across clinical groups of healthy and type I diabetes mellitus. Only *N*-acyl lipids with p-values of 0.05 or less are shown. Healthy, *n* = 52; Diabetes (type 1), *n* = 44. (**H,I**) *N*-acyl lipids annotated from a public dataset (GNPS/MassIVE: <u>MSV000084322</u>, <u>MSV000084463</u>) of (**H**) skin swabs and (**I**) soil samples of a human cadaver decomposition study.^35^ The parallel coordinates plots show the mean of the *N*-acyl lipids peak areas obtained for the different headgroups in each of the stages of decomposition. Each line represents a *N*-acyl lipid match. (**J,K**) Peak area abundances of *N*-acyl lipids annotated in public datasets from (**J**) skin (GNPS/MassIVE: <u>MSV000084322</u>) and **b)** soil (GNPS/MassIVE: <u>MSV000084463</u>) samples across different stages of decomposition of human bodies.^35^ Skin: Day0, *n* = 36; Early, *n* = 171; Active, *n* = 292; Advanced, *n* = 249. Soil: Day0, *n* = 36; Early, *n* = 171; Active, *n* = 299; Advanced, *n* = 252. (**L,M**) Peak area abundances of *N*-acyl lipids annotated in a public dataset (GNPS/MassIVE: <u>MSV000080918</u>)^30^ from mice fecal samples of mice subjected to different diets (**L**) and treatment with a cocktail of antibiotics (**M**). Antibiotics: No, *n* = 310; Yes, *n* = 27. Diet: HFD, *n* = 310; NC, *n* = 114. For the antibiotics plot, only mice fed with HFD were considered. All boxplots indicate the first (lower), median, and third (upper) quartiles, while whiskers are 1.5 times the interquartile range. Significance was tested in cases where two groups were compared using the non-parametric two-sided Mann-Whitney U test, while for more than two groups the non-parametric Kruskal-Wallis test was used, and p-values were corrected for multiple comparisons using the Benjamini-Hochberg correction. Compounds with p-values below 0.05 are highlighted in red. Icons were obtained from Bioicons.com.

**Supplementary Figure 4. MS/MS and retention time matching of *N*-acyl lipids in samples from the microbial monocultures and from the body decomposition study, related to Figure 2**. (**A**) MS/MS mirror plots and retention time matches to *N*-acyl lipids obtained via combinatorial synthesis. MS/MS spectra on the top (black) represent spectra detected in the microbial monocultures experiment (**Supplementary Figure 3F**). An unusual series of *N*-acyl 2-phenethylamines was observed and confirmed in level 1 annotation^27,28^ in two different chromatographic methods: LC1 (**A**) and LC2 (**B**) - see **Methods**. Chromatographic traces represent the exported ion chromatograms for each compound (black: sample; green: standard). (**C**) MS/MS mirror plots and retention time matches to *N*-acyl lipids obtained via combinatorial synthesis. MS/MS spectra on the top (black) represent spectra detected in the body decomposition study (**Supplementary Figure 3H-K**). Chromatographic traces represent the exported ion chromatograms for each compound (black: sample; green: standard) in two different chromatographic methods: LC1 (**C**) and LC2 (**D**) - see **Methods**. MS/MS mirror plots can be interactively inspected in the Metabolomics Spectrum Resolver^82^ with the information provided in **Supplementary Table S2**.

**Supplementary Figure 5. *N*-acyl lipids associated with HIV status, HIV plasma viral load, and neurocognitive impairment status, related to Figure 3**. (**A**) Peak area abundances of N-acyl histamines in people with HIV (PWH) and people without HIV (PWoH) (PWH, *n* = 228; PWoH, *n* = 93). (**B**) Molecular network obtained for histamine *N*-acyl lipids. (**C**) Peak area abundances of *N*-acyl polyamines in cognitively impaired and normal participants (impaired, *n* = 151; unimpaired, *n* = 162) of the HNRC. (**D**) Molecular network obtained for *N*-acyl cadaverines. Boxplots indicate the first (lower), median, and third (upper) quartiles, while whiskers are 1.5 times the interquartile range. Significance was tested using the non-parametric two-sided Mann-Whitney U test. The p-values shown are nominal p-values, and the adjusted ones (for multiple comparisons using Benjamini-Hochberg) are also available in **Supplementary Table S3**. The molecular networks were created using the Feature-Based Molecular Networking workflow^83^ within the GNPS environment^21^. The nodes are annotated based on spectral similarity matches with the *N*-acyl lipids library created. The nodes represent each MS/MS spectrum, while the edges connecting them represent their spectral similarity (threshold set to cosine > 0.7). Pie charts indicate the relative abundance of ion features in each group highlighted. This dataset is publicly available in GNPS/MassIVE under the accession number <u>MSV000092833</u>. (**E**) Bar plots showing the correlation coefficients for the association between HIV RNA viral load and various *N*-acyl lipids in the PWH (n = 203). Red bars represent positive correlations, while blue bars represent negative correlations, as determined by linear regression models. The p-values shown are nominal; adjusted p-values (corrected for multiple comparisons using the Benjamini-Hochberg method) are available in **Supplementary Table S3**. (**F**) MS/MS mirror plots and retention time matches to the pure *N*-acyl lipids standards. MS/MS spectra on the top (black) represent the ones detected in the HNRC fecal samples, while the MS/MS on the bottom (green) are the ones obtained from the standards. Chromatographic traces represent the exported ion chromatograms for each compound (black: sample; green: standard). The chromatographic method LC1 (see **Methods**) was used. MS/MS mirror plots can be interactively inspected in the Metabolomics Spectrum Resolver^82^ with the information provided in **Supplementary Table S3**. (**G**) Chromatographic traces represent the exported ion chromatograms for each compound (black: sample; green: standard), with data acquired in a different chromatographic method: LC2 (see **Methods**).

## STAR Methods

### Resource availability

#### Lead contact

Further queries and reagent requests may be directed and will be fulfilled by the lead contact, Pieter C. Dorrestein.

#### Materials availability

This study did not generate new unique reagents. All the reagents in this study were included in the key resources table.

#### Data and code availability

The *N*-acyl lipids library is available as part of the GNPS public spectral libraries (https://gnps.ucsd.edu/ProteoSAFe/gnpslibrary.jsp?library=GNPS-N-ACYL-LIPIDS-MASSQL), and is also archived in Zenodo (https://doi.org/10.5281/zenodo.14015198). All the scripts used to perform the data analyses and generate the figures are available at https://github.com/helenamrusso/N-acyl_lipids. NMR data of the pure *N*-acyl lipids has been archived at Zenodo (https://doi.org/10.5281/zenodo.14015081). All the untargeted metabolomics LC-MS/MS data are deposited on GNPS/MassIVE and publicly available under the following accession numbers: MSV000088040 (monocolonized germ-free mice), MSV000082261 (diabetes), MSV000084322 (body decomposition, skin) and MSV000084463 (body decomposition, soil), diet and antibiotics treatment (MSV000080918), MSV000095648 (microbial monocultures), MSV000092833 (HIV study), MSV000095423 (retention time and MS/MS matching), and MSV000096049 (method validation and quantification). Due to human volunteer projection constraints, the sample metadata for the HNRC cohort will be provided upon request to HNRC: https://hnrp.hivresearch.ucsd.edu/index.php/hnrc-home.

### Experimental model and study participant details

All procedures involving human participants adhered to the ethical standards established by the institutional and/or national research committee (UCSD Human Research Protections Program, HNRC IRB#172092).

## Method details

### Searches in LIPID MAPS

An initial search in LIPID MAPS^10^ was performed searching for any compound in the database that would have an amide bound, which resulted in 5,648 substructures. Ceramides were filtered out, resulting in a total of 1,240 compounds that comprised a mixture of fatty acid-derived *N*-acyl lipids, bile acid amidates, lipids attached to CoA, and *N*-acylated molecules, such as deferioxamine and other natural products. These compounds were that were manually inspected to determine if these were *N*-acyl lipids (**Supplementary Table S1**). The headgroups and acyl chain lengths were plotted (**Supplementary Figure 1A,B**) using the “seaborn.barplot” package (version 0.12.2) in Python (version 3.7.6).

### Design of the MassQL queries and creation of the N-acyl lipids library

*N*-acyl lipids were searched in the GNPS/MassIVE public datasets, which consist of ∼1.2 billion spectra, and 2,706 datasets when this search was performed in 2023. This search was initially conducted with Orbitrap public data deposited in the GNPS/MassIVE repository using the Mass Spec Query Language (MassQL)^20^. MassQL enables the filtering of public mass spectrometry data to retrieve all MS/MS spectra that contain defined and recognizable data patterns, and can also be done at the repository level. Since *N*-acyl lipids ionize well in positive ionization mode and more than 90% of the public mass spectrometry data was acquired in this mode, only positive ionization data was mined from the public domain.

The queries were developed for biologically relevant molecules conjugated to an acyl lipid chain via an amide bond (**Figure 1A**). The headgroups contained a primary or secondary amine. In total, queries were designed for 64 heads, including proteinogenic amino acids, polyamines, endocrine hormones, neurotransmitters, and other selected diagnostic-relevant metabolites, ranging from serotonin to kynurenine, and from thyroxine (T4) to glutathione (see **Supplementary Table S1** for the complete list of molecules for which a query was developed). We directed our searches to compounds in which the acyl chain varies from two up to 30 carbons, and from zero up to four unsaturations. In addition, no attachments to the acyl chain (such as hydroxylations) were considered. The queries were designed by manually inspecting reference MS/MS spectra of compounds previously synthesized,^14^ and we observed that in the vast majority of the cases, the acylium ion—resulting from the stable fatty acid chain fragmentation—was generally absent or present at very low abundances (see **Supplementary Figure 1C**). Consequently, each query was designed to target key MS/MS fragments of the headgroup, in addition to the theoretical precursor ion for each potential *N*-acyl lipid considering the precursor ions as protonated molecules ([M+H]+) (**Supplementary Figure 1D**). For the compounds for which there were no reference MS/MS libraries, the fragmentation pattern of the heads alone was manually inspected and these patterns were used in addition to the precursor ion inclusion list for each head.

Once the candidate queries were formulated, their selectivity was initially evaluated by conducting the queries in the GNPS public spectral libraries which contained 587,917 spectra of a wide variety of classes of compounds. All job links are available in **Supplementary Table S1**). A false discovery rate (FDR) was estimated by checking the retrieved spectra for each query that matched. We assessed this manually by examining the structures of the spectral matches and determining if they made sense with respect to the expected fragmentation. In cases such as alanine versus sarcosine, which are isomers and have very similar MS/MS spectra, both were accepted as true positives. Matches where the headgroup aligned with the expected fragmentation pattern were considered positive matches, while anything that did not match the headgroup was considered a false positive. Some matches were relative to putative spectra created through the propagation of a molecular network and could match related molecules but be derived from different headgroups (**Supplementary Table S1**).^84^ Therefore, hits against suspect spectra were not considered. If a high FDR was obtained for the GNPS library searches, the query was iteratively refined until the lowest possible FDR was obtained.

The final queries were then run against publicly available Orbitrap data in GNPS/MassIVE between July and October 2023. (**Supplementary Table S1**). As an additional performance assessment, but now of the repository-scale query results, all of the MS/MS spectra derived from each MassQL query were searched against all publicly available reference spectra in GNPS. The parameters used for the searches were as follows: cosine threshold as above 0.7, minimum matched fragments as 6, and precursor and fragment ion mass tolerance set to 0.02 Da. For almost all queries, the mismatched spectra comprised less than 1% of the spectral matches, except for glycine (FDR 1.5%), which had false positives to ethanolamine conjugates. Some other queries showed an FDR greater than 1%, but these false positives were matches to suspect spectra (**Supplementary Table S1**).^84^

The queries resulted in the retrieval of a total of 176,732 MS/MS spectra from Orbitrap data in the public domain (**Supplementary Figure 1E**). Six headgroups— selenomethionine, 3-methoxytyrosine, 3-iodothyronamine, levothyroxine (T4), liothyronine (T3), and homocysteine/homocystine—had no candidate *N*-acyl lipid MS/MS matches retrieved. Identical MS/MS spectra obtained from the MassQL queries were merged using MScluster^85^ to reduce duplications, keeping also only the ones that were retrieved at least twice (all GNPS job links are available in **Supplementary Table S1**). This step resulted in 1,474 unique candidate *N*-acyl lipids (**Supplementary Figure 1E**).

Because some headgroups would result in very few diagnostic ions (e.g., glycine) and could result in spurious matches, an additional filtering step was applied in these results to increase the confidence of the library. This filter was based on cosine similarity calculation between the MS/MS spectra retrieved from the queries and the reference spectra of the compounds previously synthesized.^14^ In cases where there were no reference MS/MS spectra available, a modified cosine similarity calculation was performed between the MS/MS retrieved from the query and the reference MS/MS of the headgroup. Therefore, spectra would only pass the filter if the cosine or modified cosine similarity scores would reach values above 0.7. The resulting list of MassQL-filtered spectra represented 851 unique candidate *N*-acyl amides (**Supplementary Table S1**, **Supplementary Figure 1E**), which were used to generate an MS/MS spectral library and leveraged for downstream analyses. This library, named GNPS-N-ACYL-LIPIDS-MASSQL, is currently part of the public GNPS spectral libraries and can be browsed on the web interface at the following link: https://gnps.ucsd.edu/ProteoSAFe/gnpslibrary.jsp?library=GNPS-N-ACYL-LIPIDS-MASSQL.

### N-acyl lipids repository-scale search

To expand our investigations beyond Orbitrap datasets, we conducted a repository-scale search using a fast MASST (FASST) search,^29^ an updated and faster version of the Mass Spectrometry Search Tool (MASST),^18^ against all the public domain data that were indexed in GNPS.^21^ This search is based on the cosine similarity of the queried spectra against the ones from the public domain in GNPS/MassIVE, regardless of the instrument used for data acquisition. The MGF files generated with MScluster were used as input in the FASST search pipeline, and the parameters used for this search were: cosine similarity above 0.7, minimum matched fragments as 4, and precursor ion and fragment ion tolerances as 0.02 Da. These searches were conducted using the REST web API (https://zenodo.org/records/7828220) in October 2023, using *gnps_index*. In addition to getting a table with all the MS/MS spectra matches from the public datasets, outputs from domain-specific MASSTs are also generated with this search: (1) microbeMASST: merges the FASST spectral matches against a curated database of more than 60,000 LC-MS/MS files of microbial monocultures that were taxonomically defined;^15^ (2) plantMASST: merges the FASST matches against 19,075 LC-MS/MS files of plant extracts of taxonomically defined plants;^17^ and (3) foodMASST: merges FASST matches against ∼3,500 LC-MS/MS files of foods and beverages categorized within a food ontology, collected as part of the Global FoodOmics project.^16,86^ These domain-specific MASSTs generated output tables that contain spectral matches to specific data files in the public domain that can be mapped to the curated list of taxonomy/ontology-defined metadata. Therefore, it is possible to map in which microbes, plants, or foods a particular *N*-acyl lipid was previously acquired and deposited in GNPS/MassIVE.

The FASST search enabled us to retrieve 356,542 MS/MS spectra from 61,833 files in 950 datasets and emphasizes how common *N*-acyl lipids are detected in untargeted metabolomics experiments (**Supplementary Table S1**). To explore their distribution in different organisms, tissues, and biofluids, we merged the FASST output with the ReDU (Reanalysis of Data User Interface) controlled vocabulary metadata, which enables us to do comparisons across datasets.^19^ This merged table was filtered to contain only rows relative to human (“9606|Homo sapiens”) or rodent-related datasets (“10088|Mus”, “10090|Mus musculus”, “10105|Mus minutoides”, “10114|Rattus”, “10116|Rattus norvegicus”) in the NCBITaxonomy column. Therefore, the total number of unique MS/MS spectra obtained for humans, rodents, microbes, plants, and foods was 39,525, 28,497, 29,105, 3,754, and 6,537, respectively (**Supplementary Figure 1E**).

The FASST searches are performed on indexed spectra, which are MS/MS spectra that are filtered to allow the repository searches to be conducted in seconds. To increase the level of confidence of these matches, an additional cosine similarity was performed on the FASST results to calculate the cosine similarity between the queried spectra and the pre-indexed unfiltered spectra. We filtered these results by removing any MS/MS that resulted in a cosine below 0.7 (**Supplementary Figure 1E**). Therefore, the final numbers based on ReDU and domain-specific MASST analysis were the following: 31,299 of the MS/MS spectra could be linked to human samples, 21,866 were found in rodent-related datasets, 22,589 found in microbial monocultures, 2,931 in plant-related data, and 5,576 MS/MS in foods (**Supplementary Figure 1E**).

The *N*-acyl lipids results obtained from the microbeMASST results were further filtered to: (1) remove any *N*-acyl lipid that was observed in human cell lines (which are also included in microbeMASST as control of host-derived compounds) by filtering outputs in which the taxaname was “Homo sapiens”; (2) remove *N*-acyl lipids that were retrieved only one or two times in the microbeMASST searches; and (3) remove N-acyl lipids that appear more than 5% in blanks or QCs (also added in microbeMASST). For the N-acyl lipids results of plantMASST, filters (2) and (3) were applied in a similar way. For the foodMASST results, filter (3) only was applied as there are no blanks or QC samples part of foodMASST.

The results of the queries (**Figure 1B**), body part distributions (**Figure 1C,D, Supplementary Figure 2C,D**), and microbeMASST (**Figure 2A,B**) are shown in the form of heatmaps that were created using the “seaborn.clustermap” package (version 0.12.2) in Python (version 3.7.6). The microbial classes were organized in taxonomic order according to NCBI Taxonomy ID. The UpSet plots (**Figure 2C, Supplementary Figure 3A-D**) were generated in R (https://www.R-project.org/, version 4.0.0) using the “UpSetR” package (version 1.4.0).^87^ Barplots (**Figure 2B,D**) were created using the “seaborn.barplot” package (version 0.12.2) in Python (version 3.7.6).

### Reanalysis of public data from GNPS/MassIVE

The *N*-acyl lipids library created was used to reanalyze several public datasets available in GNPS/MassIVE: (1) a study on monocolonized germ-free mice (<u>MSV000088040</u>, small intestine and colon samples); (2) a type-1 diabetes study (<u>MSV000082261</u>, urine samples); (3) a study on body decomposition (<u>MSV000084322</u> and <u>MSV000084463</u>, skin and soil samples); and (4) a dataset on the effect of diet and antibiotics consumption in colorectal cancer (<u>MSV000080918</u>, fecal samples). For each dataset, the files were downloaded from GNPS/MassIVE and processed in MZmine3 (version 3.9.0).^88^ The parameters used for each study are available in **Supplementary Table S2**, and the MZmine3 batch files are available at https://github.com/helenamrusso/N-acyl_lipids. The output files generated at the processing step (.csv file with peak areas and .mgf files with MS/MS information of each feature) were used as inputs in the Feature-Based Molecular Networking^83^ workflow in GNPS2 and wan against the *N*-acyl lipids library.^21^ The parameters of this workflow were the same for all the datasets, in which the precursor and fragment ions tolerances were set to 0.02 Da, the minimum matching peak between the queried spectra and the library was set to 4, a cosine threshold of 0.7 was applied, and no filters considered. The GNPS2 FBMN jobs can be accessed at:

- Monocolonized germ-free mice dataset (MSV000088040): https://gnps2.org/status?task=55122ffb51ab4957be51b0073abc8f04
- Diabetes dataset (MSV000082261): https://gnps2.org/status?task=ed2d2cd94179481d818356271bd7762f
- Body decomposition datasets (MSV000084322 and MSV000084463): https://gnps2.org/status?task=5f30dc8527bb456190daf2e772bf399d
- Impact of diet and antibiotics consumption in colorectal cancer dataset (<u>MSV000080918</u>): https://gnps2.org/status?task=ba8fd7da3d654d1d949d4beb398b4152

For the monocolonized germ-free mice, a heatmap was obtained to show the log2 of the fold change of *N*-acyl lipids levels of colonized and monocolonized mice in relation to the germ-free group (**Supplementary Figure 3E**). To achieve this, the median of each feature annotated as an *N*-acyl lipid was calculated and the log2(FC) was calculated. The small intestine and colon samples were considered separately. The heatmap was created using the “seaborn.clustermap” package (version 0.12.2) in Python (version 3.7.6). The microbial strains were organized in taxonomic order according to NCBI Taxonomy ID and their classes were also mapped to the heatmap. The *N*-acyl lipids were organized in ascending order in the number of carbons and unsaturations.

To determine if the *N*-acyl lipid matches had a significant correlation with any of the groups in each study, the peak areas (exported .csv file from MZmine3) of the matching compounds in the datasets were plotted as boxplots using the “seaborn.boxplot” package (version 0.12.2) in Python (version 3.7.6).

Line plots were obtained for the body decomposition dataset to show the changes in *N*-acyl lipids with increasing body decomposition levels (**Supplementary Figure 3H,I**). The mean of each feature annotated as an *N*-acyl lipid was calculated for each stage of decomposition and a line plot was obtained using Matplotlib (version 3.2.1) in Python (version 3.7.6). Finally, volcano plots were obtained for the diet and antibiotics treatment study, where the log2(FC) was calculated for each *N*-acyl lipid (**Figure 2E,F**), and scatter plots were obtained with Matplotlib (version 3.2.1) in Python (version 3.7.6).

### Bacterial cultures screening

#### Bacterial strains, growth conditions, and metabolite extraction

The bacteria used in this study are listed in the **Supplementary Table S2**. All bacteria cultures were started from glycerol stock and incubated at 37°C anaerobically (10% CO_2_, 7.5% H_2_, 82.5% N_2_) in a filtered BHI medium (**Supplementary Table S2**) at a pH adjusted to 7.2 using 5 N NaOH. Cultures were normalized at OD_600_ = 0.02 before being diluted 1/10 in 1 mL of mPYG medium (**Supplementary Table S2**) and incubated for 48-72 h at 37°C in a 2 mL deep-well plate. Following bacterial growth, 400 µL of culture was transferred to a new 2 mL deep-well plate and extracted overnight at 4°C with 1.2 mL of pre-chilled 50% MeOH/H_2_O. Samples were dried in a CentriVap and stored at -80°C until LC-MS/MS analysis.

#### LC-MS/MS untargeted metabolomics analysis

Samples were resuspended in 200 µL of 50% MeOH/H_2_O with 1 µM of sulfadimethoxine as internal standard, incubated at -20°C overnight before being centrifuged at 21,130 x g. Then, 150 µL was transferred into a 2 mL glass vial containing an insert. The chromatographic separation was performed by reversed-phase polar C18 (Kinetex Polar C18, 100 mm x 2.1 mm, 2,6 µm, 100 A pore size with a guard column, Phenomenex) using a Vanquish UHPLC system coupled to a Q-Exactive Orbitrap mass spectrometer (Thermo Fisher Scientific). The mobile phase consisted of solvent A (water + 0.1% formic acid) and solvent B (ACN + 0.1% formic acid) and the column compartment was kept at 40 °C. Five microlitres of the samples were injected and eluted at a flow rate of 0.5 mL/min using the following gradient: 0 – 1.1 min 5% B, 1.1 – 7.5 min 40% B, 7.5 – 8.5 min 99% B, 8.5 – 9.5 min 99% B, 9.5 – 10 min 5% B, 10 – 10.5 min 5% B, 10.5 – 10.75 min 99% B, 10.75 – 11.25 min 99% B, 11.5 – 12 min 5% B. Mass spectrometry (MS) analysis was performed using electrospray ionization (ESI) in positive ionization mode. The parameters were set as follows: Sheath gas flow 53 L/min, auxiliary gas flow rate 14 L/min, sweep gas flow 3 L/min, spray voltage 3.5 kV, inlet capillary to 269°C, and auxiliary gas heater set to 438°C. MS scan range 100 – 1000 *m/z* with a resolution (R*_m/z_* _200_) of 35,000 with 1 microscans. The automatic gain control (AGC) target was set to 1E6 with a maximum injection time of 100 ms. Up to 5 MS/MS spectra per MS1 were collected with a resolution (R*_m/z_* _200_) set to 17,500 with 1 microscans, maximum injection time of 150 ms with an AGC target of 5E5. The isolation window was set to 1 *m/z* and the isolation offset at 0 *m/z*. The normalized collision energy was acquired with an increase stepwise at 25, 40, and 60. The apex trigger was set to 2 – 15 s and a dynamic exclusion of 5 s. Isotopes were excluded from the analysis. The data is publicly available on GNPS/MassIVE <u>MSV000095648</u>.

#### Data processing, molecular networking, and data visualization

The LC-MS/MS files were converted to .mzML using MSConvert (ProteoWizard, Palo Alto, CA, USA)^89^ and processed in MZmine4 (version 4.0.8).^88^ The parameters used for data processing are listed in **Supplementary Table S2**. The exported files were subjected to the FBMN^83^ workflow in GNPS2.^21^ The parameters used for the library search for *N*-acyl lipids annotation were as follows: precursor and fragment ion tolerances set to 0.02 Da, 4 minimum matched peaks between the queried spectra and the library, a cosine threshold of 0.7, and no filters considered. The GNPS2 FBMN job can be accessed at https://gnps2.org/status?task=cff8c1a5895b4b9b917c36ad9444c635.

A heatmap was obtained to show the variations of the features annotated as *N*-acyl lipids with regard to the microbial media. A median value was obtained for all the media samples, and for all the bacteria. A filter was applied to only consider the features that were detected in both microbial replicates. Therefore, if there were any zero values for one of the microbial replicates, all values were set to zero. The log2(FC) was calculated and plotted using the “seaborn.clustermap” package (version 0.12.2) in Python (version 3.7.6). The microbial strains were organized in taxonomic order according to NCBI Taxonomy ID and their classes were also mapped to the heatmap. The *N*-acyl lipids were organized in ascending order in the number of carbons and unsaturations.

### Combinatorial reactions of N-acyl lipids

Fatty acid (1 eq.) and 2 mL of THF were added to a 20 mL scintillation vial with a magnetic stir bar. To this solution, solid EDC (1 eq.) and neat DIPEA (1 eq.) were subsequently added, and the solution was stirred at RT. After 15 minutes, amine compound (1 eq.) in 1mL H_2_O was added, and the reaction was stirred for 14 h. To proceed with LC-MS/MS analyses, 1 µL of the reaction mixture was diluted in 1 mL of LC-MS grade MeOH.

### HIV Neurobehavioral Research Center (HNRC) cohort study

#### Cohort - clinical ratings

The neurocognitive test battery included measures that assess seven neurocognitive domains commonly affected by HIV, including verbal fluency, speeded information processing, executive functions, learning, memory, working memory, and motor.^90^ Raw scores for each test were converted to demographically corrected T-scores and used to derive global clinical ratings using a published standardized algorithm that classifies the presence and severity of NCI. Global ratings have a nine-point scale ranging from normal (1-4), to mildly impaired (5-6), to moderately or severely impaired (7-9), with a clinical rating of greater than or equal to 5 indicative of NCI.^91^ This is consistent with Frascati’s Criteria for diagnosing HIV-associated neurocognitive disorder, which requires at least mild neurocognitive impairment in at least two of the seven neurocognitive ability domains.^92^

#### Sample preparation

The study was conducted following protocols approved by the UCSD Human Research Protections Program (https://irb.ucsd.edu/), and all participants gave written informed consent before participating. Stool samples were processed using a recently developed automated pipeline designed for simultaneous extraction of metagenomic and metabolomic data.^93^ For the metabolites extraction, the swabs were placed into Matrix Tubes (ThermoFisher Scientific, MA, USA) containing 400 μL of 95% ethanol (v/v), and the tubes were sealed using the Capit-All automated capping instrument (ThermoFisher Scientific, MA, USA). The tubes were shaken at 1,200 rpm for 2 min using a SpexMiniG plate shaker, followed by centrifugation at 2,700 g for 5 min. Part of the supernatant (200 µL) was transferred to a deep well plate with an 8-channel pipette, dried down in a vacuum centrifuge concentrator at room temperature for approximately 5 h, and stored at -80°C until LC-MS/MS analyses.

#### LC-MS/MS untargeted metabolomics analysis

Prior to the analyses, the samples were resuspended in 200 µL of acetonitrile:H_2_O (1:1, v/v) with 100 μg/L sulfadimethoxine as the internal standard, sonicated for 10 min, centrifuged at 450 g for 10 min, and 150 µL of the supernatant was transferred to a shallow 96 well plate for analyses. The extracts were injected (5 µL) into a Vanquish UHPLC system coupled to a Q-Exactive Orbitrap mass spectrometer (Thermo Fisher Scientific). The chromatographic separation was achieved by reverse-phase polar C18 (150 × 2.1 mm, 2.6 μm particle size, 100 A pore size; Phenomenex, Torrance) with a SecurityGuard C18 column (2.1 mm ID) at 30 °C column temperature. The mobile phase consisted of solvents A (water) and B (ACN) both containing 0.1% formic acid, and the flow rate was set at 0.5 mL/min. The following gradient was applied: 0-1 min 5% B, 1-7 min 5-99% B, 7-8 min 99% B, 8-8.5 min 99-5% B, 8-10 min 5%B. Mass spectrometry (MS) analysis was performed using electrospray ionization (ESI) in positive ionization mode, and the parameters were set as follows: sheath gas flow 53 L/min, auxiliary gas flow rate 14 L/min, sweep gas flow 3 L/min, spray voltage 3.5 kV, inlet capillary to 269°C, and auxiliary gas heater 400 °C. MS1 scan range was set to *m/z* 100-1500 with a resolution (R*_m/z_* _200_) of 35,000, automatic gain control (AGC) target as 5.0E4, and maximum injection time of 100 ms. Up to 5 MS/MS spectra per MS1 were collected with a resolution (R*_m/z_* _200_) set to 17,500, AGC target as 5.0E4, and maximum injection time of 100 ms. The isolation window was set to 3 *m/z* and the isolation offset was set to 0.5 *m/z*. The normalized collision energy was acquired with an increased stepwise from 20 to 30 to 40%. The apex trigger was set to 2 - 15 s, the minimum AGC target for the MS/MS spectrum was 5.0E3, and a dynamic precursor exclusion of 10 s was selected. This dataset is publicly available in GNPS/MassIVE under the following accession number:

#### Data processing and Feature-Based Molecular Networking

The LC-MS/MS files were converted to .mzML using MSConvert (ProteoWizard, Palo Alto, CA, USA)^89^ and processed in MZmine3 (version 3.9.0).^88^ The parameters used for data processing are listed in **Supplementary Table S2**. The exported files were subjected to the FBMN^83^ workflow in GNPS2.^21^ No filters were applied to the data, and the precursor and MS/MS fragment ion tolerances were set to 0.02 Da. A molecular network was created, in which the edges were filtered to have a cosine score above 0.7 and at least four matched fragments. Similarly, the parameters for the *N*-acyl lipids library search were set to have a cosine value above 0.7 and at least four matched fragments. The GNPS2 FBMN job can be accessed at https://gnps2.org/status?task=ee34ee95908749dd81ee9a62fbdac98e. The molecular networks were visualized in Cytoscape^94^ (version 3.10.0).

#### Metagenomic data sequencing

Fecal samples were processed for shotgun metagenomics sequencing as previously described.^95^ The metagenomic libraries were normalized by iSeq (Illumina) read count distribution to generate a final pool that made sequencing on the NovaSeq more efficient.^96^ Raw sequence files were demultiplexed using BaseSpace (Illumina, CA, USA), and quality-filtered following a previous protocol.^97^ The filtered reads were aligned to the Web of Life database^98^ using bowtie2,^99^, and the settings used were the following: maximum and minimum mismatch penalties (mp=[1,1]), a penalty for ambiguities (np=1; default), read and reference gap open- and extend penalties (rdg=[0,1], rfg=[0,1]), a minimum alignment score for an alignment to be considered valid (score-min=[L,0,-0.05]), a defined number of distinct, valid alignments (k=16), and the suppression of SAM records for unaligned reads, as well as SAM headers (no-unal, no-hd). A feature table was obtained by converting the resulting alignments using the Web of Life Toolkit App.^100^

#### Metagenomic data processing

The metagenomic data processing was performed as previously described.^101^ The sequence data were filtered for all adapters known to fastp (version 0.23.4) in paired-end mode by explicitly specifying a known adapters file.^102^ Fastp also removed sequences shorter than 45 nucleotides with -l, a flag to filter the minimum length of each sequence. Each sample was then filtered against each genome in the human pangenome,^103^ as well as both T2T-CHM13v2.0^104^ and GRCh38,^105^ using minimap2^106^ (version 2.26-r1175) with “-ax sr” for short read mode. The data were first run in paired-end mode, and then run in single-end mode, per genome. Each successive run was converted from SAM to FASTQ using samtools^107^ (version 1.17) with arguments -f 12 -F 256 -N for paired-end data and -f 4 -F 256 for single-end. The single-end data are repaired using fastq_pair^108^ (version 1.0) specifying a table size of 50M with -t. Compute support was provided with GNU Parallel^109^ (version 20180222). Single-end FASTQ output from samtools was split into R1 and R2 with a custom Rust program, with rust-bio for parsing^110^ (version 1.4.0). Data were multiplexed with sed and demultiplexed using a custom Python script. Shotgun sequencing data were then uploaded to and processed through Qiita^111^ (Study ID 11135). Sequence adapter and host filtering were executed using qp-fastp-minimap2 version 2022.04. Subsequently, Woltka^100^ version 0.1.4 (qp-woltka 2022.09) with the Web of Life 2 database was employed for taxonomic and functional predictions. Genomic coverages were computed, and features with less than 25% coverage were excluded.^112^ To further enhance data quality, a prevalence filter using QIIME 2 v2023.5^113^ was applied, eliminating features present in less than 10% of samples to mitigate the inclusion of erroneous and low-quality reads. The resulting feature table was utilized for downstream analysis.

#### Microbe-metabolite multi-omics associations

Co-occurrence probabilities between microbes and metabolites were calculated using mmvec^40^ (version 1.0.4) as a Qiime2^113^ plugin. Mmvec takes as input the relative abundance matrix for the sequencing data and the feature abundance table for the ion features, and through a neural networking approach, conditional probabilities of observing molecules based on the abundance of each microbe are calculated. A subset of samples with both metabolite and microbiome data were used for this analysis (225 samples). The mmvec parameters were as follows: -p-batch-size 50, -p-num-testing-examples 5, -p-epochs 50, -p-learning-rate 1e-4. All other parameters for the analyses were set as the default values. EMPeror^114^ was used to visually inspect the feature-feature biplots (https://view.qiime2.org/). The spheres on the plot were colored based on which group (HIV+ vs HIV-) the molecules were most abundant, and the arrows indicate the 30 most important OTUs retrieved from the analyses (*i.e.,* higher magnitude of the vector using Euclidean distance from the origin) (**Figure 3D**). The co-occurrence probabilities were also inspected at the microbial order taxonomic level for the histamine and cadaverine *N*- acyl lipids. Only histamine-C2:0 and histamine-C3:0 had co-occurrence probabilities > 6.0, and a network was obtained for the microbial orders that were shared between both compounds (**Figure 3E**). The network was visualized in Cytoscape^94^ (version 3.10.0). All inputs and outputs from mmvec and the Cytoscape visualization file are available at https://github.com/helenamrusso/N-acyl_lipids.

#### Microbial cultures from the multi-omics analysis

*Holdemanella biformis* DSM 3989, *Catenibacterium mitsuokai* DSM 15897, *Megasphaera* sp. DSMZ 102144, *Dorea longicatena* DSM 13814, *Prevotella buccae* D17, *Eubacterium siraeum* DSM 15702*, Collinsella aerofaciens* ATCC 25986*, Roseburia inulinivorans* DSM 16841, and *Streptococcus thermophilus* LMD-9 were selected for microbial culturing based on the multi-omics results and strains availability. these microbes were cultured in 200 µL in BHI medium (**Supplementary Table S2**) for 72 h at 37°C in an anaerobic chamber supplemented with 100 µM of cadaverine, putrescine, and histamine. Samples were extracted overnight at 4°C using 600 µL of pre-chilled 50% MeOH/H_2_O. Samples were then dried using a CentriVap and stored at -80°C until resuspension.

#### Untargeted LC-MS/MS analysis of microbes from the multi-omics HIV analysis

The microbial extracts were resuspended in H_2_O (100%) containing 1 µM of sulfamethazine to achieve a concentration of 50 mg/mL, incubated at -20°C overnight, and centrifuged at 21,130 x g. Then, 120 µL of the solution was transferred to a 2 mL glass vial containing an insert for LC-MS/MS analysis. The samples were injected (2 µL) into a Vanquish UHPLC system coupled to a Q-Exactive Orbitrap mass spectrometer (Thermo Fisher Scientific). The chromatographic separation was achieved by reverse-phase polar C18 (Kinetex Polar C18, 100 × 2.1 mm, 2.6 μm particle size, 100 A pore size; Phenomenex, Torrance) with a SecurityGuard C18 column (2.1 mm ID) at 40 °C column temperature. The mobile phase consisted of solvents A (water) and B (ACN) both containing 0.1% formic acid, and the flow rate was set at 0.5 mL/min. The gradient employed consisted of 0-1 min 1% B, 1-7.5 min 5-99% B, 7.5-9.3 min 99% B, 9.3-9.5 min 99-1% B, 9.5-11 min 1%B. Mass spectrometry (MS) analysis was performed using electrospray ionization (ESI) in positive ionization mode, and the parameters were set as follows: sheath gas flow 53 L/min, auxiliary gas flow rate 14 L/min, sweep gas flow 3 L/min, spray voltage 3.5 kV, inlet capillary to 269°C, and auxiliary gas heater 430 °C. MS1 scan range was set to *m/z* 100-1500 with a resolution (R*_m/z_* _200_) of 35,000, automatic gain control (AGC) target as 5.0E4, and maximum injection time of 100 ms. Up to 5 MS/MS spectra per MS1 were collected with a resolution (R*_m/z_* _200_) set to 17,500, AGC target as 5.0E5, and maximum injection time of 50 ms. The isolation window was set to 2 *m/z* and the isolation offset was set to 0 *m/z*. The normalized collision energy was acquired with an increased stepwise from 25 to 40 to 60%. The apex trigger was set to 1 to 5 s, the minimum AGC target for the MS/MS spectrum was 8.0E3, and a dynamic precursor exclusion of 10 s was selected. The data was deposited in GNPS/MassIVE and is publicly available at <u>MSV000095648</u>.

## Retention time and MS/MS matching with combinatorial synthetic standard reaction mixtures

Extracts from skin samples of the body decomposition study (<u>MSV000084322</u>) and the HIV study (<u>MSV000092833</u>) were available in our laboratory for additional analyses to get retention time and MS/MS spectral matching between synthetic standards and biological samples. In addition, the samples from the microbial monocultures described in the “*Bacterial cultures screening*” section were used to confirm the microbial production of selected *N*-acyl lipids. Therefore, the biological samples and the synthetic standards were subjected to LC-MS/MS analyses. The dried extracts were resuspended in 150 µL of MeOH:H_2_O (1:1, v/v) for the microbial extracts (n = 2) and the body decomposition samples (n = 4), while the HIV samples (n = 4) were resuspended in 150 µL of H_2_O (100%). The same method described in “*Untargeted LC-MS/MS analysis of microbes from the multi-omics HIV analysis*” was used to acquire the data. However, two different gradients were used to evaluate the retention time matching between the synthetic *N*-acyl lipids and the compounds present in the biological samples: the first gradient (LC1) consisted of 0-1 min 1% B, 1-7.5 min 5-99% B, 7.5-9.3 min 99% B, 9.3-9.5 min 99-1% B, 9.5-11 min 1%B; and the second gradient (LC2) consisted of 0-1.5 min 1% B, 1.5-10.5 min 5-99% B, 10.5-12.3 min 99% B, 12.3-12.5 min 99-1% B, 12.5-14 min 1%B. The acquired LC-MS/MS data was deposited in GNPS/MassIVE and is publicly available at <u>MSV000095423</u>.

### Obtention of pure N-acyl lipids

Pure *N*-acyl lipids were acquired commercially from Sigma-Aldrich, Aldlab Chemicals, or EnamineStore. More specifically, N-(2-(1H-imidazol-4-yl)ethyl)acetamide (histamine-C2:0, purity 98%), *N*-(5-aminopentyl)acetamide (cadaverine-C2:0, purity 95%), *N*-(3,4-dihydroxyphenethyl)acetamide (dopamine-C2:0, purity 95%), and *N*-(2-(5-hydroxy-1H-indol-3-yl)ethyl)acetamide (serotonin-C2:0, purity >99%) were acquired from Sigma-Aldrich; *N*-(2-(1H-imidazol-4-yl)ethyl)propionamide (histamine-C3:0, purity 98%), *N*-(2-(1H-imidazol-4-yl)ethyl)butyramide (histamine-C4:0, purity 95%), *N*-(2-(1H-imidazol-4-yl)ethyl)pentanamide (histamine-C5:0, purity 98%), and *N*-(5-aminopentyl)propionamide (cadaverine-C3:0, purity 95%) were acquired from EnamineStore; and *N*-(2-(1H-imidazol-4-yl)ethyl)hexanamide (histamine-C6:0, purity 95%), *N*-(5-aminopentyl)pentanamide (cadaverine-C5:0, purity 95%), *N*-(5-aminopentyl)hexanamide (cadaverine-C6:0, purity 95%), *N*-(5-aminopentyl)heptanamide (cadaverine-C7:0, purity 95%), and propionyl-L-tryptophan (tryptophan-C3:0, purity 95%) were acquired from Aldlab Chemicals.

The structure of these *N*-acyl lipids was confirmed by NMR ^1^H. NMR spectra were collected at 298 K on a 600 MHz Bruker Avance III spectrometer fitted with a 1.7 mm triple resonance cryoprobe with z-axis gradients. The spectra were acquired in CD_3_OD-*d_4_* or CDCl_3_-*d_1_*, which was chosen based on the solubility of the compounds. The shifts are reported in ppm and calibrated against the residual solvent signals at δ_H_ 3.31 and 7.26 for CD_3_OD-*d_4_* and CDCl_3_-*d_1_* respectively. The deuterated solvents were acquired from Cambridge Isotope Laboratories, Inc. (Andover, USA). The NMR data acquired were deposited in Zenodo (https://doi.org/10.5281/zenodo.14015081).

### Quantification of N-acyl lipids in biological samples

The LC-MS/MS method used for the analyses of the method validation and quantification was the same as previously described in the “*Retention time and MS/MS matching with combinatorial synthetic standard reaction mixtures*” section, employing gradient LC1. The analytical method was performed according to the International Conference on Harmonization (ICH) guidelines^115^ for histamine-C2:0, histamine-C3:0, histamine-C4:0, histamine-C5:0, cadaverine-C2:0, cadaverine-C3:0, cadaverine-C5:0, cadaverine-C6:0, and dopamine-C2:0. The method was validated based on the evaluation of the following parameters: specificity, precision (repeatability and intermediate precision), linearity, limit of detection (LOD), limit of quantification (LOQ), and accuracy. Detailed information regarding the methodology used for each of them is described below, and all the figures of merit are available in **Supplementary Table S3**. The validation was performed using sample P3_D9_Sample_X3157299 from the HNRC cohort that would contain the compounds of interest. Skyline^116^ (version 23.1) was used to extract the peak areas of the *N*-acyl lipids. The method employed reached the acceptance criteria specified for each parameter (**Supplementary Table S3**). For quantification in biological samples, 148 samples of the HIV cohort were available and injected in the validated method (samples were resuspended in 100 µL of H_2_O containing 1 µM of sulfamethazine). For the calculation of the amounts in the samples, it was estimated that 10 mg of stool sample would be the starting material, as previously described,^117^ and the extraction yield was also extrapolated to 100%. In addition, all the samples of the microbial monocultures described in the “*Microbial cultures from the multi-omics analysis*” were also analyzed. The injection volume was set to 2 µL for all samples.

#### Specificity

The specificity was determined by injecting a blank solution containing only the internal standard (sulfadimethazine), and an injection of a solution containing all the *N*- acyl lipids (n=3). The relative standard deviation (RSD) was calculated based on each peak’s retention time in the P3_D9_Sample_X3157299 sample. The MS and MS/MS spectra confirmed the specificity and identity of these compounds. The retention times of the peaks of interest were as follows: histamine-C2:0, 0.58 min; histamine-C3:0, 0.73 min; histamine-C4:0, 1.13 min; histamine-C5:0, 2.29 min; cadaverine-C2:0, 0.61 min; cadaverine-C3:0, 0.78 min; cadaverine-C5:0, 2.47 min; cadaverine-C6:0, 2.94 min; dopamine-C2:0, 2.60 min. These compounds didn’t show interferences compared to the solution containing only the mixture of standards.

#### Precision (repeatability and intermediate precision)

The precision of the method was determined by analyzing the P3_D9_Sample_X3157299 sample in six replicates (n=6), and the repeatability (intra-day precision) was estimated as the RSD of the standards concentrations (µg/mL) measured in two consecutive days. The concentrations calculated for the compounds on both days are available in **Supplementary Table S3**. The RSD values were lower than 5%, and the F-test between the two days showed no significant difference at F=0.05.

#### Linearity

The linearity of the method was determined by calibration curves in concentration ranges comprising each compound at the samples of interest. A stock solution containing 60 µg/mL of each *N*-acyl lipid was prepared in H_2_O (100%) and used to acquire calibration curves for all the compounds simultaneously. From this solution, 6 to 13 points were prepared with levels ranging from 0.001 to 20 µg/mL, and each concentration level was injected in triplicate. The analytical curves were built based on the nominal concentrations, and the average between the ratios of each compound and the internal standard used (Ratio = A_compound_/A_IS_). A polynomial equation was obtained for each curve, and the correlation coefficients (R) were calculated for each compound. The linear ranges and R coefficients are available in **Supplementary Table S3**.

#### Limit of detection and limit of quantification

LODs and LOQs were estimated by the mean of the slopes (a) and the standard deviation of the y-intercept (Sb) on three calibration curves (linear regression was used) in three low concentrations for each compound (0.002 to 0.02 µg/mL). A linear regression was used in this estimation. These limits were calculated by the following equations: LOD = (3.3*Sb)/a and LOQ = (10*Sb)/a. All the slopes, intercepts, LODs, and LOQs are shown in **Supplementary Table S3**.

#### Accuracy

The accuracy of the method was determined by recovery analyses. For this, known amounts of the solution containing the standards were spiked to the P3_E10_Sample_x3137731 and P3_G2_Sample_X3148765 sample solutions in two different concentrations (low and high) considering the predetermined calibration curve and concentration range. Three replicates for each level were injected and analyzed in the validated method. The accuracy was determined by the difference between the theoretical and experimental concentration values and the values were within the acceptance range of 80–120%.

### Statistical analyses

Statistical tests were performed using the non-parametric Mann-Whitney U test in cases where two groups were being compared (diabetes, diet, and antibiotic treatment - **Supplementary Figure 3G,L,M, Figure 2E,F**), or with the non-parametric Kruskal-Wallis for more than two groups (body decomposition - **Supplementary Figure 3J,K**). The p-values were corrected for multiple comparisons using the Benjamini-Hochberg correction. The statistical tests were done with the “scipy.stats” package (version 1.7.3), and the p-values corrections with the “statsmodels.stats.multitest” (version 0.11.1) in Python (version 3.7.6).

For the HNRC study, the differences in individual /V-acyl lipids between the study groups were compared using a multivariate linear mixed-effects model with fixed covariates for HIV status (PWH vs. PWoH) and neurocognitive impairment status (impaired vs. unimpaired) (∼ HIV status + neurocognitive impairment), while accounting for random effects within individual samples (∼1 | Subject) using the MaAsLin2 package in R (version 4.2.1). Lipid values were log-transformed, and zero values were imputed with half the minimum value prior to analysis. The regression coefficients from the linear model were illustrated as a forest plot using the ’ggplot2’ (version 3.5.1) package in R (version 4.2.1). To visualize the correlation coefficients from the linear model with only fixed effects (i.e., the association between CD4/CD8 ratio or plasma viral load), a horizontal bar plot was created using ’ggplot2’ (version 3.5.1). The color palettes were selected from the RColorBrewer(version 1.1.3) package in R (version 4.2.1).

## Supplementary table titles and legends

**Supplementary Table S1.** Queries jobs, queries results, and body part distribution for rodents and humans, related do Figure 1.

**Supplementary Table S2.** *N*-acyl lipids chain length diversity, evidence of microbial N-acyl lipids, and reanalysis of public datasets, related to Figure 2.

**Supplementary Table S3.** HIV and neurocognition study, multiomics results and quantification of N-acyl lipids, related to Figure 3.

**Supplementary Table S4.** Microbial production and activity data of *N*-acyl lipids, related to Figure 4.

## References

1. Chang, F.-Y., Siuti, P., Laurent, S., Williams, T., Glassey, E., Sailer, A.W., Gordon, D.B., Hemmerle, H., and Voigt, C.A. (2021). Gut-inhabiting Clostridia build human GPCR ligands by conjugating neurotransmitters with diet- and human-derived fatty acids. Nat Microbiol 6, 792–805.

2. Mann, A., Smoum, R., Trembovler, V., Alexandrovich, A., Breuer, A., Mechoulam, R., and Shohami, E. (2015). Palmitoyl Serine: An Endogenous Neuroprotective Endocannabinoid-Like Entity After Traumatic Brain Injury. J. Neuroimmune Pharmacol. 10, 356–363.

3. Waluk, D.P., Vielfort, K., Derakhshan, S., Aro, H., and Hunt, M.C. (2013). N-Acyl taurines trigger insulin secretion by increasing calcium flux in pancreatic β-cells. Biochem. Biophys. Res. Commun. 430, 54–59.

4. Aichler, M., Borgmann, D., Krumsiek, J., Buck, A., MacDonald, P.E., Fox, J.E.M., Lyon, J., Light, P.E., Keipert, S., Jastroch, M., et al. (2017). N-acyl Taurines and Acylcarnitines Cause an Imbalance in Insulin Synthesis and Secretion Provoking β Cell Dysfunction in Type 2 Diabetes. Cell Metab. 25, 1334–1347.e4.

5. Arul Prakash, S., and Kamlekar, R.K. (2021). Function and therapeutic potential of N-acyl amino acids. Chem. Phys. Lipids 239, 105114.

6. Long, J.Z., Svensson, K.J., Bateman, L.A., Lin, H., Kamenecka, T., Lokurkar, I.A., Lou, J., Rao, R.R., Chang, M.R., Jedrychowski, M.P., et al. (2016). The secreted enzyme PM20D1 regulates lipidated amino acid uncouplers of mitochondria. Cell 166, 424–435.

7. Connor, M., Vaughan, C.W., and Vandenberg, R.J. (2010). N-acyl amino acids and N-acyl neurotransmitter conjugates: neuromodulators and probes for new drug targets. Br. J. Pharmacol. 160, 1857–1871.

8. Jörgensen, A.M., Wibel, R., and Bernkop-Schnürch, A. (2023). Biodegradable cationic and ionizable cationic lipids: A roadmap for safer pharmaceutical excipients. Small 19, e2206968.

9. Mokhtari, V., Afsharian, P., Shahhoseini, M., Kalantar, S.M., and Moini, A. (2017). A review on various uses of N-acetyl cysteine. Cell J. 19, 11–17.

10. Conroy, M.J., Andrews, R.M., Andrews, S., Cockayne, L., Dennis, E.A., Fahy, E., Gaud, C., Griffiths, W.J., Jukes, G., Kolchin, M., et al. (2024). LIPID MAPS: update to databases and tools for the lipidomics community. Nucleic Acids Res. 52, D1677–D1682.

11. Xue, J., Chi, L., Tu, P., Lai, Y., Liu, C.-W., Ru, H., and Lu, K. (2021). Detection of gut microbiota and pathogen produced N-acyl homoserine in host circulation and tissues. NPJ Biofilms Microbiomes 7, 53.

12. Tan, B., O’Dell, D.K., Yu, Y.W., Monn, M.F., Hughes, H.V., Burstein, S., and Walker, J.M. (2010). Identification of endogenous acyl amino acids based on a targeted lipidomics approach. J. Lipid Res. 51, 112–119.

13. Wood, P.L. (2019). Targeted lipidomics and metabolomics evaluations of cortical neuronal stress in schizophrenia. Schizophr. Res. 212, 107–112.

14. Gentry, E.C., Collins, S.L., Panitchpakdi, M., Belda-Ferre, P., Stewart, A.K., Carrillo Terrazas, M., Lu, H.-H., Zuffa, S., Yan, T., Avila-Pacheco, J., et al. (2023). Reverse metabolomics for the discovery of chemical structures from humans. Nature. 10.1038/s41586-023-06906-8.

15. Zuffa, S., Schmid, R., Bauermeister, A., P Gomes, P.W., Caraballo-Rodriguez, A.M., El Abiead, Y., Aron, A.T., Gentry, E.C., Zemlin, J., Meehan, M.J., et al. (2024). microbeMASST: a taxonomically informed mass spectrometry search tool for microbial metabolomics data. Nat Microbiol 9, 336–345.

16. West, K.A., Schmid, R., Gauglitz, J.M., Wang, M., and Dorrestein, P.C. (2022). foodMASST a mass spectrometry search tool for foods and beverages. NPJ Sci Food 6, 22.

17. Gomes, P.W.P., Mannochio-Russo, H., Schmid, R., Zuffa, S., Damiani, T., Quiros-Guerrero, L.-M., Caraballo-Rodríguez, A.M., Zhao, H.N., Yang, H., Xing, S., et al. (2024). plantMASST - Community-driven chemotaxonomic digitization of plants. bioRxiv. 10.1101/2024.05.13.593988.

18. Wang, M., Jarmusch, A.K., Vargas, F., Aksenov, A.A., Gauglitz, J.M., Weldon, K., Petras, D., da Silva, R., Quinn, R., Melnik, A.V., et al. (2020). Mass spectrometry searches using MASST. Nat. Biotechnol. 38, 23–26.

19. Jarmusch, A.K., Wang, M., Aceves, C.M., Advani, R.S., Aguirre, S., Aksenov, A.A., Aleti, G., Aron, A.T., Bauermeister, A., Bolleddu, S., et al. (2020). ReDU: a framework to find and reanalyze public mass spectrometry data. Nat. Methods 17, 901–904.

20. Jarmusch, A.K., Aron, A.T., Petras, D., Phelan, V.V., Bittremieux, W., Acharya, D.D., Ahmed, M.M.A., Bauermeister, A., Bertin, M.J., Boudreau, P.D., et al. (2022). A Universal Language for Finding Mass Spectrometry Data Patterns. bioRxiv, 2022.08.06.503000. 10.1101/2022.08.06.503000.

21. Wang, M., Carver, J.J., Phelan, V.V., Sanchez, L.M., Garg, N., Peng, Y., Nguyen, D.D., Watrous, J., Kapono, C.A., Luzzatto-Knaan, T., et al. (2016). Sharing and community curation of mass spectrometry data with Global Natural Products Social Molecular Networking. Nat. Biotechnol. 34, 828–837.

22. Mohanty, I., Mannochio-Russo, H., Schweer, J.V., El Abiead, Y., Bittremieux, W., Xing, S., Schmid, R., Zuffa, S., Vasquez, F., Muti, V.B., et al. (2024). The underappreciated diversity of bile acid modifications. Cell 187, 1801–1818.e20.

23. Watanabe, K., Yasugi, E., and Oshima, M. (2000). How to search the glycolipid data in “LIPIDBANK for web” the newly developed lipid database in japan. Trends Glycosci. Glycotechnol. 12, 175–184.

24. Aimo, L., Liechti, R., Hyka-Nouspikel, N., Niknejad, A., Gleizes, A., Götz, L., Kuznetsov, D., David, F.P.A., van der Goot, F.G., Riezman, H., et al. (2015). The SwissLipids knowledgebase for lipid biology. Bioinformatics 31, 2860–2866.

25. Bhandari, S., Bisht, K.S., and Merkler, D.J. (2021). The Biosynthesis and Metabolism of the N-Acylated Aromatic Amino Acids: N-Acylphenylalanine, N-Acyltyrosine, N-Acyltryptophan, and N-Acylhistidine. Front Mol Biosci 8, 801749.

26. Ghosh, A.K., and Shahabi, D. (2021). Synthesis of amide derivatives for electron deficient amines and functionalized carboxylic acids using EDC and DMAP and a catalytic amount of HOBt as the coupling reagents. Tetrahedron Lett. 63. 10.1016/j.tetlet.2020.152719.

27. Sumner, L.W., Amberg, A., Barrett, D., Beale, M.H., Beger, R., Daykin, C.A., Fan, T.W.-M., Fiehn, O., Goodacre, R., Griffin, J.L., et al. (2007). Proposed minimum reporting standards for chemical analysis Chemical Analysis Working Group (CAWG) Metabolomics Standards Initiative (MSI). Metabolomics 3, 211–221.

28. Schymanski, E.L., Jeon, J., Gulde, R., Fenner, K., Ruff, M., Singer, H.P., and Hollender, J. (2014). Identifying small molecules via high resolution mass spectrometry: communicating confidence. Environ. Sci. Technol. 48, 2097–2098.

29. Batsoyol, N., Pullman, B., Wang, M., Bandeira, N., and Swanson, S. (2022). P-Massive: A Real-Time Search Engine for a Multi-Terabyte Mass Spectrometry Database. In SC22: International Conference for High Performance Computing, Networking, Storage and Analysis, pp. 1–15.

30. Shalapour, S., Lin, X.-J., Bastian, I.N., Brain, J., Burt, A.D., Aksenov, A.A., Vrbanac, A.F., Li, W., Perkins, A., Matsutani, T., et al. (2017). Inflammation-induced IgA+ cells dismantle anti-liver cancer immunity. Nature 551, 340–345.

31. Song, X., Zhang, H., Zhang, Y., Goh, B., Bao, B., Mello, S.S., Sun, X., Zheng, W., Gazzaniga, F.S., Wu, M., et al. (2023). Gut microbial fatty acid isomerization modulates intraepithelial T cells. Nature 619, 837–843.

32. Wu, M., Zheng, W., Song, X., Bao, B., Wang, Y., Ramanan, D., Yang, D., Liu, R., Macbeth, J.C., Do, E.A., et al. (2024). Gut complement induced by the microbiota combats pathogens and spares commensals. Cell 187, 897–913.e18.

33. Cheng, J., Venkatesh, S., Ke, K., Barratt, M.J., and Gordon, J.I. (2024). A human gut *Faecalibacterium prausnitzii* fatty acid amide hydrolase. Science 386. 10.1126/science.ado6828.

34. Cheng, A.G., Ho, P.-Y., Aranda-Díaz, A., Jain, S., Yu, F.B., Meng, X., Wang, M., Iakiviak, M., Nagashima, K., Zhao, A., et al. (2022). Design, construction, and in vivo augmentation of a complex gut microbiome. Cell 185, 3617–3636.e19.

35. Burcham, Z.M., Belk, A.D., McGivern, B.B., Bouslimani, A., Ghadermazi, P., Martino, C., Shenhav, L., Zhang, A.R., Shi, P., Emmons, A., et al. (2024). A conserved interdomain microbial network underpins cadaver decomposition despite environmental variables. Nat Microbiol 9, 595–613.

36. Watrous, J., Roach, P., Alexandrov, T., Heath, B.S., Yang, J.Y., Kersten, R.D., van der Voort, M., Pogliano, K., Gross, H., Raaijmakers, J.M., et al. (2012). Mass spectral molecular networking of living microbial colonies. Proc. Natl. Acad. Sci. U. S. A. 109, E1743–E1752.

37. Quinn, R.A., Melnik, A.V., Vrbanac, A., Fu, T., Patras, K.A., Christy, M.P., Bodai, Z., Belda-Ferre, P., Tripathi, A., Chung, L.K., et al. (2020). Global chemical effects of the microbiome include new bile-acid conjugations. Nature 579, 123–129.

38. Quinn, R.A., Nothias, L.-F., Vining, O., Meehan, M., Esquenazi, E., and Dorrestein, P.C. (2017). Molecular networking as a drug discovery, drug metabolism, and precision medicine strategy. Trends Pharmacol. Sci. 38, 143–154.

39. Buggert, M., Frederiksen, J., Noyan, K., Svärd, J., Barqasho, B., Sönnerborg, A., Lund, O., Nowak, P., and Karlsson, A.C. (2014). Multiparametric bioinformatics distinguish the CD4/CD8 ratio as a suitable laboratory predictor of combined T cell pathogenesis in HIV infection. J. Immunol. 192, 2099–2108.

40. Morton, J.T., Aksenov, A.A., Nothias, L.F., Foulds, J.R., Quinn, R.A., Badri, M.H., Swenson, T.L., Van Goethem, M.W., Northen, T.R., Vazquez-Baeza, Y., et al. (2019). Learning representations of microbe-metabolite interactions. Nat. Methods 16, 1306–1314.

41. Mann, E.R., Lam, Y.K., and Uhlig, H.H. (2024). Short-chain fatty acids: linking diet, the microbiome and immunity. Nat. Rev. Immunol. 24, 577–595.

42. Silva, Y.P., Bernardi, A., and Frozza, R.L. (2020). The role of short-chain fatty acids from gut Microbiota in gut-brain communication. Front. Endocrinol. 11, 25.

43. van der Hee, B., and Wells, J.M. (2021). Microbial regulation of host physiology by short-chain fatty acids. Trends Microbiol. 29, 700–712.

44. Verma, A., Bhagchandani, T., Rai, A., Nikita, Sardarni, U.K., Bhavesh, N.S., Gulati, S., Malik, R., and Tandon, R. (2024). Short-chain fatty acid (SCFA) as a connecting link between Microbiota and gut-lung axis-A potential therapeutic intervention to improve lung health. ACS Omega 9, 14648–14671.

45. González-Hernández, L.A., Ruiz-Briseño, M.D.R., Sánchez-Reyes, K., Alvarez-Zavala, M., Vega-Magaña, N., López-Iñiguez, A., Díaz-Ramos, J.A., Martínez-Ayala, P., Soria-Rodriguez, R.A., Ramos-Solano, M., et al. (2019). Alterations in bacterial communities, SCFA and biomarkers in an elderly HIV-positive and HIV-negative population in western Mexico. BMC Infect. Dis. 19, 234.

46. Enriquez, A.B., Ten Caten, F., Ghneim, K., Sekaly, R.-P., and Sharma, A.A. (2023). Regulation of immune homeostasis, inflammation, and HIV persistence by the microbiome, short-chain fatty acids, and bile acids. Annu. Rev. Virol. 10, 397–422.

47. Hollenbaugh, J.A., Munger, J., and Kim, B. (2011). Metabolite profiles of human immunodeficiency virus infected CD4+ T cells and macrophages using LC-MS/MS analysis. Virology 415, 153–159.

48. Mahalingam, S.S., Jayaraman, S., Bhaskaran, N., Schneider, E., Faddoul, F., Paes da Silva, A., Lederman, M.M., Asaad, R., Adkins-Travis, K., Shriver, L.P., et al. (2023). Polyamine metabolism impacts T cell dysfunction in the oral mucosa of people living with HIV. Nat. Commun. 14, 399.

49. Moon, J.-Y., Zolnik, C.P., Wang, Z., Qiu, Y., Usyk, M., Wang, T., Kizer, J.R., Landay, A.L., Kurland, I.J., Anastos, K., et al. (2018). Gut microbiota and plasma metabolites associated with diabetes in women with, or at high risk for, HIV infection. EBioMedicine 37, 392–400.

50. Li, X., Wu, T., Jiang, Y., Zhang, Z., Han, X., Geng, W., Ding, H., Kang, J., Wang, Q., and Shang, H. (2018). Plasma metabolic changes in Chinese HIV-infected patients receiving lopinavir/ritonavir based treatment: Implications for HIV precision therapy. Cytokine 110, 204–212.

51. Ding, Y., Lin, H., Chen, X., Zhu, B., Xu, X., Xu, X., Shen, W., Gao, M., and He, N. (2021). Comprehensive metabolomics profiling reveals common metabolic alterations underlying the four major non-communicable diseases in treated HIV infection. EBioMedicine 71, 103548.

52. Cassol, E., Misra, V., Holman, A., Kamat, A., Morgello, S., and Gabuzda, D. (2013). Plasma metabolomics identifies lipid abnormalities linked to markers of inflammation, microbial translocation, and hepatic function in HIV patients receiving protease inhibitors. BMC Infect. Dis. 13, 203.

53. Babu, H., Sperk, M., Ambikan, A.T., Rachel, G., Viswanathan, V.K., Tripathy, S.P., Nowak, P., Hanna, L.E., and Neogi, U. (2019). Plasma metabolic signature and abnormalities in HIV-infected individuals on long-term successful antiretroviral therapy. Metabolites 9, 210.

54. Taylor, B.C., Weldon, K.C., Ellis, R.J., Franklin, D., Groth, T., Gentry, E.C., Tripathi, A., McDonald, D., Humphrey, G., Bryant, M., et al. (2020). Depression in individuals coinfected with HIV and HCV is associated with systematic differences in the gut microbiome and metabolome. mSystems 5. 10.1128/mSystems.00465-20.

55. Pedersen, M., Nielsen, C.M., and Permin, H. (1991). HIV antigen-induced release of histamine from basophils from HIV infected patients. Mechanism and relation to disease progression and immunodeficiency. Allergy 46, 206–212.

56. Armstrong, A.J.S., Shaffer, M., Nusbacher, N.M., Griesmer, C., Fiorillo, S., Schneider, J.M., Preston Neff, C., Li, S.X., Fontenot, A.P., Campbell, T., et al. (2018). An exploration of Prevotella-rich microbiomes in HIV and men who have sex with men. Microbiome 6, 198.

57. Fusco, W., Lorenzo, M.B., Cintoni, M., Porcari, S., Rinninella, E., Kaitsas, F., Lener, E., Mele, M.C., Gasbarrini, A., Collado, M.C., et al. (2023). Short-chain fatty-acid-producing bacteria: Key components of the human gut Microbiota. Nutrients 15. 10.3390/nu15092211.

58. Hövelmann, Y., Steinert, K., Hübner, F., and Humpf, H.-U. (2020). Identification of a novel N-caprylhistamine-β-glucoside from tomato fruits and LC-MS/MS-based food screening for imidazole alkaloids. Food Chem. 312, 126068.

59. Hövelmann, Y., Hahn, M., Hübner, F., and Humpf, H.-U. (2019). Detection of novel cytotoxic imidazole alkaloids in tomato products by LC-MS/MS. J. Agric. Food Chem. 67, 3670–3678.

60. Takao, K., Noguchi, K., Hashimoto, Y., Shirahata, A., and Sugita, Y. (2015). Synthesis and evaluation of fatty acid amides on the N-oleoylethanolamide-like activation of peroxisome proliferator activated receptor α. Chem. Pharm. Bull. 63, 278–285.

61. Huang, W., Rha, G.B., Han, M.-J., Eum, S.Y., András, I.E., Zhong, Y., Hennig, B., and Toborek, M. (2008). PPARalpha and PPARgamma effectively protect against HIV-induced inflammatory responses in brain endothelial cells. J. Neurochem. 107, 497–509.

62. Crakes, K.R., Santos Rocha, C., Grishina, I., Hirao, L.A., Napoli, E., Gaulke, C.A., Fenton, A., Datta, S., Arredondo, J., Marco, M.L., et al. (2019). PPARα-targeted mitochondrial bioenergetics mediate repair of intestinal barriers at the host-microbe intersection during SIV infection. Proc. Natl. Acad. Sci. U. S. A. 116, 24819–24829.

63. Roan, N.R., and Greene, W.C. (2007). A seminal finding for understanding HIV transmission. Cell 131, 1044–1046.

64. Herbert, C., Luies, L., Loots, D.T., and Williams, A.A. (2023). The metabolic consequences of HIV/TB co-infection. BMC Infect. Dis. 23, 536.

65. Zhang, Y., Xie, Z., Zhou, J., Li, Y., Ning, C., Su, Q., Ye, L., Ai, S., Lai, J., Pan, P., et al. (2022). The altered metabolites contributed by dysbiosis of gut microbiota are associated with microbial translocation and immune activation during HIV infection. Front. Immunol. 13, 1020822.

66. Wu, R., Chen, X., Kang, S., Wang, T., Gnanaprakasam, J.R., Yao, Y., Liu, L., Fan, G., Burns, M.R., and Wang, R. (2020). De novo synthesis and salvage pathway coordinately regulate polyamine homeostasis and determine T cell proliferation and function. Sci. Adv. 6, eabc4275.

67. Puleston, D.J., Baixauli, F., Sanin, D.E., Edwards-Hicks, J., Villa, M., Kabat, A.M., Kamiński, M.M., Stanckzak, M., Weiss, H.J., Grzes, K.M., et al. (2021). Polyamine metabolism is a central determinant of helper T cell lineage fidelity. Cell 184, 4186– 4202.e20.

68. Lee, S.H., Kim, S.O., Lee, H.D., and Chung, B.C. (1998). Estrogens and polyamines in breast cancer: their profiles and values in disease staging. Cancer Lett. 133, 47–56.

69. Paik, M.-J., Lee, S., Cho, K.-H., and Kim, K.-R. (2006). Urinary polyamines and N-acetylated polyamines in four patients with Alzheimer’s disease as their N-ethoxycarbonyl-N-pentafluoropropionyl derivatives by gas chromatography-mass spectrometry in selected ion monitoring mode. Anal. Chim. Acta 576, 55–60.

70. Kovács, T., Mikó, E., Vida, A., Sebő, É., Toth, J., Csonka, T., Boratkó, A., Ujlaki, G., Lente, G., Kovács, P., et al. (2019). Cadaverine, a metabolite of the microbiome, reduces breast cancer aggressiveness through trace amino acid receptors. Sci. Rep. 9, 1300.

71. Salzman, S.K., and Stepita-Klauco, M. (1981). Inhibition of evoked dopamine release by monopropionylcadaverine in vitro. Pharmacol. Biochem. Behav. 15, 119–123.

72. Murray, K.E., Shaw, K.J., Adams, R.F., and Conway, P.L. (1993). Presence of N-acyl and acetoxy derivatives of putrescine and cadaverine in the human gut. Gut 34, 489–493.

73. Salzman, S.K., and Stepita-Klauco, M. (1981). Cadaverine in the rat brain: regional distribution and acylation of [14C]cadaverine in vivo and uptake in vitro. J. Neurochem. 37, 1308–1315.

74. Mayers, J.R., Varon, J., Zhou, R.R., Daniel-Ivad, M., Beaulieu, C., Bhosle, A., Glasser, N.R., Lichtenauer, F.M., Ng, J., Vera, M.P., et al. (2024). A metabolomics pipeline highlights microbial metabolism in bloodstream infections. Cell 187, 4095– 4112.e21.

75. Dudkina, N., Park, H.B., Song, D., Jain, A., Khan, S.A., Flavell, R.A., Johnson, C.H., Palm, N.W., and Crawford, J.M. (2024). Human AKR1C3 binds agonists of GPR84 and participates in an expanded polyamine pathway. Cell Chem. Biol. 10.1016/j.chembiol.2024.07.011.

76. Elmassry, M.M., Sugihara, K., Chankhamjon, P., Camacho, F.R., Wang, S., Sugimoto, Y., Chatterjee, S., Chen, L.A., Kamada, N., and Donia, M.S. (2024). A meta-analysis of the gut microbiome in inflammatory bowel disease patients identifies disease-associated small molecules. bioRxivorg. 10.1101/2024.02.07.579278.

77. Merali, S., Barrero, C.A., Sacktor, N.C., Haughey, N.J., Datta, P.K., Langford, D., and Khalili, K. (2014). Polyamines: Predictive biomarker for HIV-associated neurocognitive disorders. J. AIDS Clin. Res. 5, 1000312.

78. Lyons, D.E., Beery, J.T., Lyons, S.A., and Taylor, S.L. (1983). Cadaverine and aminoguanidine potentiate the uptake of histamine in vitro in perfused intestinal segments of rats. Toxicol. Appl. Pharmacol. 70, 445–458.

79. Sánchez-Pérez, S., Comas-Basté, O., Costa-Catala, J., Iduriaga-Platero, I., Veciana-Nogués, M.T., Vidal-Carou, M.C., and Latorre-Moratalla, M.L. (2022). The rate of histamine degradation by diamine oxidase is compromised by other biogenic amines. Front. Nutr. 9, 897028.

80. Hui, J.Y., and Taylor, S.L. (1985). Inhibition of in vivo histamine metabolism in rats by foodborne and pharmacologic inhibitors of diamine oxidase, histamine N-methyltransferase, and monoamine oxidase. Toxicol. Appl. Pharmacol. 81, 241– 249.

81. Schmid, R., Petras, D., Nothias, L.-F., Wang, M., Aron, A.T., Jagels, A., Tsugawa, H., Rainer, J., Garcia-Aloy, M., Dührkop, K., et al. (2021). Ion identity molecular networking for mass spectrometry-based metabolomics in the GNPS environment. Nat. Commun. 12, 3832.

82. Bittremieux, W., Chen, C., Dorrestein, P.C., Schymanski, E.L., Schulze, T., Neumann, S., Meier, R., Rogers, S., and Wang, M. (2020). Universal MS/MS Visualization and Retrieval with the Metabolomics Spectrum Resolver Web Service. bioRxiv, 2020.05.09.086066. 10.1101/2020.05.09.086066.

83. Nothias, L.-F., Petras, D., Schmid, R., Dührkop, K., Rainer, J., Sarvepalli, A., Protsyuk, I., Ernst, M., Tsugawa, H., Fleischauer, M., et al. (2020). Feature-based molecular networking in the GNPS analysis environment. Nat. Methods 17, 905– 908.

84. Bittremieux, W., Avalon, N.E., Thomas, S.P., Kakhkhorov, S.A., Aksenov, A.A., Gomes, P.W.P., Aceves, C.M., Caraballo-Rodríguez, A.M., Gauglitz, J.M., Gerwick, W.H., et al. (2023). Open access repository-scale propagated nearest neighbor suspect spectral library for untargeted metabolomics. Nat. Commun. 14, 8488.

85. Frank, A.M., Monroe, M.E., Shah, A.R., Carver, J.J., Bandeira, N., Moore, R.J., Anderson, G.A., Smith, R.D., and Pevzner, P.A. (2011). Spectral archives: extending spectral libraries to analyze both identified and unidentified spectra. Nat. Methods 8, 587–591.

86. Gauglitz, J.M., West, K.A., Bittremieux, W., Williams, C.L., Weldon, K.C., Panitchpakdi, M., Di Ottavio, F., Aceves, C.M., Brown, E., Sikora, N.C., et al. (2022). Enhancing untargeted metabolomics using metadata-based source annotation. Nat. Biotechnol. 40, 1774–1779.

87. Conway, J.R., Lex, A., and Gehlenborg, N. (2017). UpSetR: an R package for the visualization of intersecting sets and their properties. Bioinformatics 33, 2938– 2940.

88. Schmid, R., Heuckeroth, S., Korf, A., Smirnov, A., Myers, O., Dyrlund, T.S., Bushuiev, R., Murray, K.J., Hoffmann, N., Lu, M., et al. (2023). Integrative analysis of multimodal mass spectrometry data in MZmine 3. Nat. Biotechnol. 10.1038/s41587-023-01690-2.

89. Kessner, D., Chambers, M., Burke, R., Agus, D., and Mallick, P. (2008). ProteoWizard: open source software for rapid proteomics tools development. Bioinformatics 24, 2534–2536.

90. Heaton, R.K., Franklin, D.R., Ellis, R.J., McCutchan, J.A., Letendre, S.L., Leblanc, S., Corkran, S.H., Duarte, N.A., Clifford, D.B., Woods, S.P., et al. (2011). HIV-associated neurocognitive disorders before and during the era of combination antiretroviral therapy: differences in rates, nature, and predictors. J. Neurovirol. 17, 3–16.

91. Woods, S.P., Rippeth, J.D., Frol, A.B., Levy, J.K., Ryan, E., Soukup, V.M., Hinkin, C.H., Lazzaretto, D., Cherner, M., Marcotte, T.D., et al. (2004). Interrater reliability of clinical ratings and neurocognitive diagnoses in HIV. J. Clin. Exp. Neuropsychol. 26, 759–778.

92. Antinori, A., Arendt, G., Becker, J.T., Brew, B.J., Byrd, D.A., Cherner, M., Clifford, D.B., Cinque, P., Epstein, L.G., Goodkin, K., et al. (2007). Updated research nosology for HIV-associated neurocognitive disorders. Neurology 69, 1789–1799.

93. Brennan, C., Belda-Ferre, P., Zuffa, S., Charron-Lamoureux, V., Mohanty, I., Ackermann, G., Allaband, C., Ambre, M., Boyer, T., Bryant, M., et al. (2024). Clearing the plate: a strategic approach to mitigate well-to-well contamination in large-scale microbiome studies. mSystems, e0098524.

94. Shannon, P., Markiel, A., Ozier, O., Baliga, N.S., Wang, J.T., Ramage, D., Amin, N., Schwikowski, B., and Ideker, T. (2003). Cytoscape: a software environment for integrated models of biomolecular interaction networks. Genome Res. 13, 2498– 2504.

95. Sanders, J.G., Nurk, S., Salido, R.A., Minich, J., Xu, Z.Z., Zhu, Q., Martino, C., Fedarko, M., Arthur, T.D., Chen, F., et al. (2019). Optimizing sequencing protocols for leaderboard metagenomics by combining long and short reads. Genome Biol. 20, 226.

96. Brennan, C., Salido, R.A., Belda-Ferre, P., Bryant, M., Cowart, C., Tiu, M.D., González, A., McDonald, D., Tribelhorn, C., Zarrinpar, A., et al. (2023). Maximizing the potential of high-throughput next-generation sequencing through precise normalization based on read count distribution. mSystems 8, e0000623.

97. Armstrong, G., Martino, C., Morris, J., Khaleghi, B., Kang, J., DeReus, J., Zhu, Q., Roush, D., McDonald, D., Gonazlez, A., et al. (2022). Swapping metagenomics preprocessing pipeline components offers speed and sensitivity increases. mSystems 7, e0137821.

98. Zhu, Q., Mai, U., Pfeiffer, W., Janssen, S., Asnicar, F., Sanders, J.G., Belda-Ferre, P., Al-Ghalith, G.A., Kopylova, E., McDonald, D., et al. (2019). Phylogenomics of 10,575 genomes reveals evolutionary proximity between domains Bacteria and Archaea. Nat. Commun. 10, 5477.

99. Langmead, B., and Salzberg, S.L. (2012). Fast gapped-read alignment with Bowtie 2. Nat. Methods 9, 357–359.

100. Zhu, Q., Huang, S., Gonzalez, A., McGrath, I., McDonald, D., Haiminen, N., Armstrong, G., Vázquez-Baeza, Y., Yu, J., Kuczynski, J., et al. (2022). Phylogeny-aware analysis of metagenome community ecology based on matched reference genomes while bypassing taxonomy. mSystems 7, e0016722.

101. Sepich-Poore, G.D., McDonald, D., Kopylova, E., Guccione, C., Zhu, Q., Austin, G., Carpenter, C., Fraraccio, S., Wandro, S., Kosciolek, T., et al. (2024). Robustness of cancer microbiome signals over a broad range of methodological variation. Oncogene 43, 1127–1148.

102. Chen, S., Zhou, Y., Chen, Y., and Gu, J. (2018). fastp: an ultra-fast all-in-one FASTQ preprocessor. Bioinformatics 34, i884–i890.

103. Liao, W.-W., Asri, M., Ebler, J., Doerr, D., Haukness, M., Hickey, G., Lu, S., Lucas, J.K., Monlong, J., Abel, H.J., et al. (2023). A draft human pangenome reference. Nature 617, 312–324.

104. Rhie, A., Nurk, S., Cechova, M., Hoyt, S.J., Taylor, D.J., Altemose, N., Hook, P.W., Koren, S., Rautiainen, M., Alexandrov, I.A., et al. (2023). The complete sequence of a human Y chromosome. Nature 621, 344–354.

105. Schneider, V.A., Graves-Lindsay, T., Howe, K., Bouk, N., Chen, H.-C., Kitts, P.A., Murphy, T.D., Pruitt, K.D., Thibaud-Nissen, F., Albracht, D., et al. (2017). Evaluation of GRCh38 and de novo haploid genome assemblies demonstrates the enduring quality of the reference assembly. Genome Res. 27, 849–864.

106. Li, H. (2021). New strategies to improve minimap2 alignment accuracy. Bioinformatics 37, 4572–4574.

107. Danecek, P., Bonfield, J.K., Liddle, J., Marshall, J., Ohan, V., Pollard, M.O., Whitwham, A., Keane, T., McCarthy, S.A., Davies, R.M., et al. (2021). Twelve years of SAMtools and BCFtools. Gigascience 10. 10.1093/gigascience/giab008.

108. Edwards, J.A., and Edwards, R.A. (2019). Fastq-pair: efficient synchronization of paired-end fastq files. 10.1101/552885.

109. Tange, O. (2018). Gnu Parallel 2018 (Zenodo).

110. Köster, J. (2016). Rust-Bio: a fast and safe bioinformatics library. Bioinformatics 32, 444–446.

111. Gonzalez, A., Navas-Molina, J.A., Kosciolek, T., McDonald, D., Vázquez-Baeza, Y., Ackermann, G., DeReus, J., Janssen, S., Swafford, A.D., Orchanian, S.B., et al. (2018). Qiita: rapid, web-enabled microbiome meta-analysis. Nat. Methods 15, 796–798.

112. Hakim, D., Wandro, S., Zengler, K., Zaramela, L.S., Nowinski, B., Swafford, A., Zhu, Q., Song, S.J., Gonzalez, A., McDonald, D., et al. (2022). Zebra: Static and dynamic genome cover thresholds with overlapping references. mSystems 7, e0075822.

113. Bolyen, E., Rideout, J.R., Dillon, M.R., Bokulich, N.A., Abnet, C.C., Al-Ghalith, G.A., Alexander, H., Alm, E.J., Arumugam, M., Asnicar, F., et al. (2019). Reproducible, interactive, scalable and extensible microbiome data science using QIIME 2. Nat. Biotechnol. 37, 852–857.

114. Vázquez-Baeza, Y., Pirrung, M., Gonzalez, A., and Knight, R. (2013). EMPeror: a tool for visualizing high-throughput microbial community data. Preprint, 10.1186/2047-217x-2-16 https://doi.org/10.1186/2047-217x-2-16.

115. ICH (2005). International Conference on Harmonization (ICH) of Technical Requirements for Registration of Pharmaceuticals for Human Use. Preprint at Topic Q9: Quality Risk Management Geneva.

116. Adams, K.J., Pratt, B., Bose, N., Dubois, L.G., St John-Williams, L., Perrott, K.M., Ky, K., Kapahi, P., Sharma, V., MacCoss, M.J., et al. (2020). Skyline for Small Molecules: A Unifying Software Package for Quantitative Metabolomics. J. Proteome Res. 19, 1447–1458.

117. Melnik, A.V., da Silva, R.R., Hyde, E.R., Aksenov, A.A., Vargas, F., Bouslimani, A., Protsyuk, I., Jarmusch, A.K., Tripathi, A., Alexandrov, T., et al. (2017). Coupling targeted and untargeted mass spectrometry for metabolome-microbiome-wide association studies of human fecal samples. Anal. Chem. 89, 7549–7559.

