## Supplementary Figures for "The microbiome diversifies *N*-acyl lipid pools - including short-chain fatty acid-derived compounds"

**Author affiliations:**
[1] Skaggs School of Pharmacy and Pharmaceutical Sciences, University of California San Diego, La Jolla, CA, USA
[2] Department of Laboratory Medicine, University of Groningen, University Medical Center Groningen, 9713 GZ Groningen, the Netherlands
[3] Turku Bioscience Center, University of Turku and Åbo Akademi University, 20520 Turku, Finland
[4] Division of Host-Microbe Systems & Therapeutics, Department of Pediatrics, University of California San Diego, La Jolla, CA 92093, USA
[5] Department of Pediatrics, University of California San Diego, La Jolla, California, USA
[6] Neurosciences Graduate Program, University of California San Diego, La Jolla, California, USA
[7] Department of Neurosciences, University of California San Diego, San Diego, CA 92093, USA
[8] Department of Psychiatry, University of California San Diego, San Diego, CA 92093, USA
[9] HIV Neurobehavioral Research Program, University of California San Diego, San Diego, CA 92093, USA
[10] Department of Medicine, University of California San Diego, La Jolla, CA, USA
[11] Department of Bioengineering, University of California, San Diego, La Jolla, CA, 92093, USA
[12] Center for Microbiome Innovation, University of California, San Diego, La Jolla, CA, 92093, USA

[13] Larsson-Rosenquist Foundation Mother-Milk-Infant Center of Research Excellence (MOMI CORE) and the Human Milk Institute (HMI), University of California San Diego, La Jolla, CA, 92093, USA
[14] Department of Immunology, Harvard Medical School, Boston, MA 02115, USA
[15] Key Laboratory of Multi-Cell Systems, Shanghai Institute of Biochemistry and Cell Biology, Center for Excellence in Molecular Cell Science, Chinese Academy of Sciences, University of Chinese Academy of Sciences, Shanghai, 200031, China
[16] Department of Microbiology, University of Tennessee, Knoxville, Tennessee, USA
[17] Department of Pediatrics, Division of Pediatric Endocrinology, University of California San Diego, California, USA
[18] Rady Children's Hospital San Diego, San Diego, California, USA
[19] Division of Pediatric Endocrinology, Children's Hospital of Orange County, Orange, CA, USA
[20] Chiba University-UC San Diego Center for Mucosal Immunology, Allergy, and Vaccines, La Jolla, California 92093, USA
[21] Department of Computer Science and Engineering, University of California Riverside, Riverside, CA, USA
[22] Department of Computer Science and Engineering, University of California, San Diego, La Jolla, CA, USA
[23] Halıcıoğlu Data Science Institute, University of California, San Diego, La Jolla, CA, USA
[24] Shu Chien-Gene Lay Department of Bioengineering, University of California, San Diego, La Jolla, CA, USA
[25] Collaborative Mass Spectrometry Innovation Center, Skaggs School of Pharmacy and Pharmaceutical Sciences, University of California San Diego, La Jolla, CA, USA
[26] Department of Pharmacology, University of California San Diego, La Jolla, CA, 92093, USA

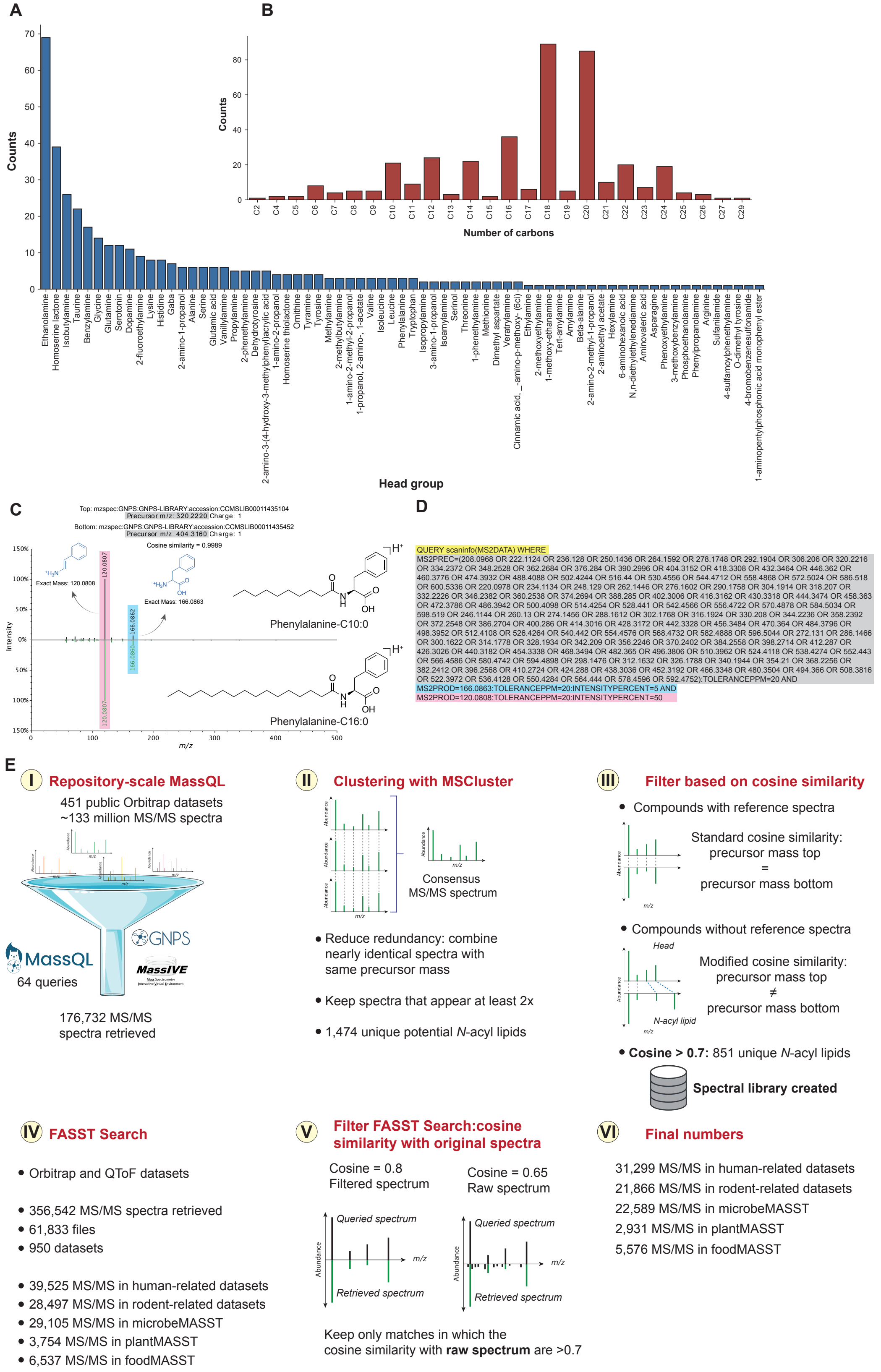


**Supplementary Figure 1. Distribution of *N*-acyl lipids in structural databases and mass spectrometry repository searches, related to Figure 1. A)** Diversity and relative frequency of *N*-acyl lipids headgroups and **(B)** lipid chain lengths documented in LIPID MAPS. This analysis excludes ceramide acylations. **C)** *N*-acyl lipid query strategy: representative MS/MS spectrum of phenylalanine-C10:0 (CCMSLIB00011435104) and phenylalanine-C16:0 (CCMSLIB00011435452). The spectra show nearly identical fragmentation patterns enabling the creation of the MassQL query to retrieve the MS/MS spectra of this family of lipids. **D)** MassQL query for phenylalanine headgroup where we initiate to return all MS/MS spectra (in yellow) that fulfill the following criteria: the precursor ion has to match one of the expected precursor *m/z* values specified (gray), as well as the most diagnostic *m/z* fragments of the head portion (blue and pink) with their indicated error tolerances and minimum relative intensities. **E)** Strategy followed to create the *N*-acyl lipids library and expand to biological interpretations. (I) MassQL queries were designed and run against the Orbitrap datasets in the GNPS/MassIVE repository. (II) The spectra were clustered using MSCluster to reduce redundancy. (III) A cosine similarity filter was applied to keep the higher confidence *N*-acyl lipids spectra. (IV) The clustered spectra were searched using FASST searches against the whole repository (including Orbitrap and QToF datasets), and human and rodent-related datasets were tagged using ReDU, and microbial, plant, and food-related datasets were also tagged using domain-specific MASSTs. (V) The spectra retrieved from the FASST searches were filtered to keep the matches in which the raw (unfiltered) spectra resulted in cosine similarity above 0.7. (VI) Summary of the results obtained with this workflow. Icons were obtained from Bioicons.com.

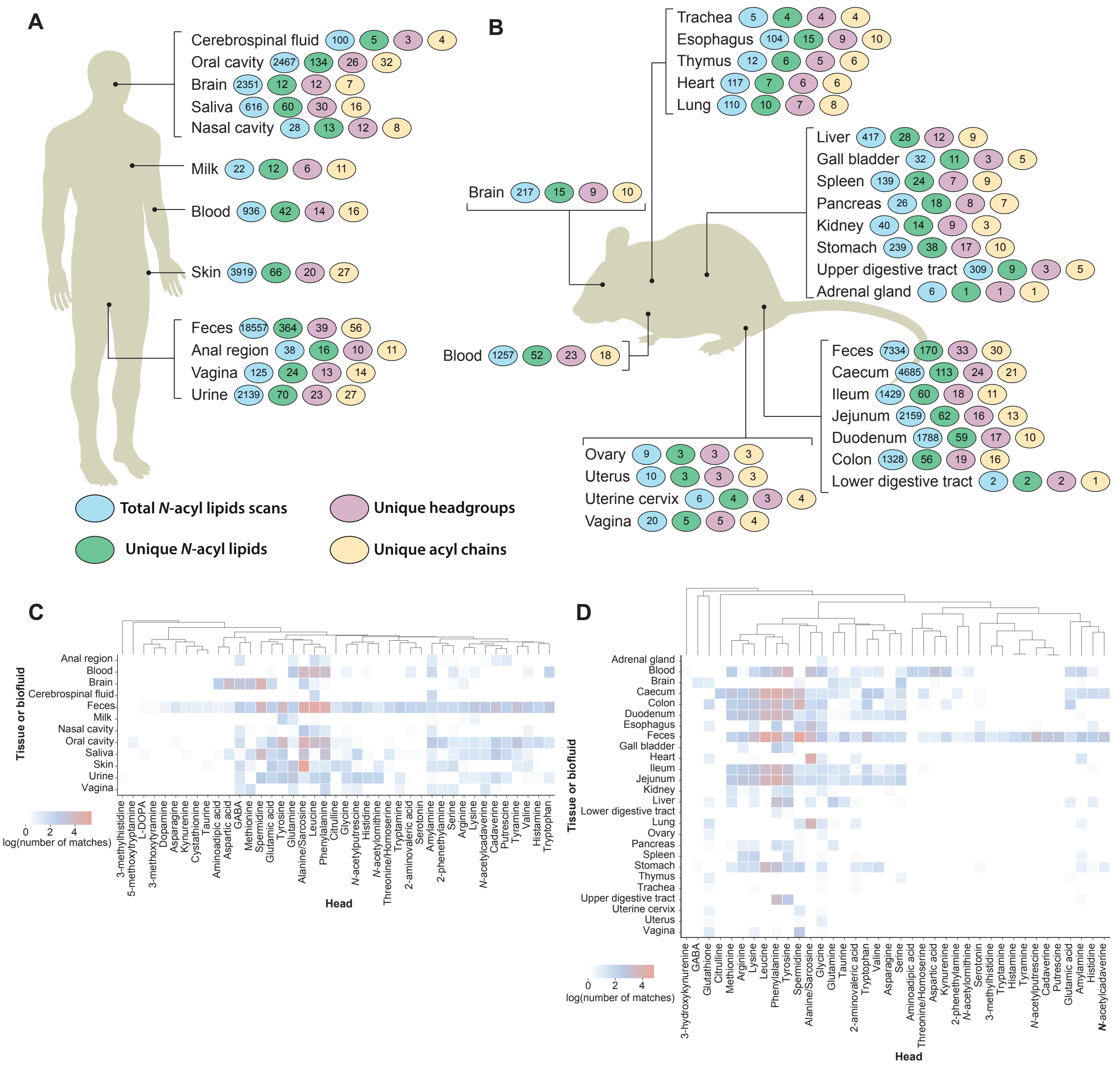


**Supplementary Figure 2. Distribution of *N*-acyl lipids obtained from FASST searches among different tissues or biofluids, related to Figure 1.** Summary of the occurrences in the public domain in (**A**) human and (**B**) rodent-related datasets. Heatmaps show the distribution of the number of matches grouped by headgroup in different tissues and biofluids with metadata available in ReDU for (**C**) human and (**D**) rodent-related public datasets. All heatmaps are shown as log values of the matches obtained from the repository. Icons were obtained from Bioicons.com.

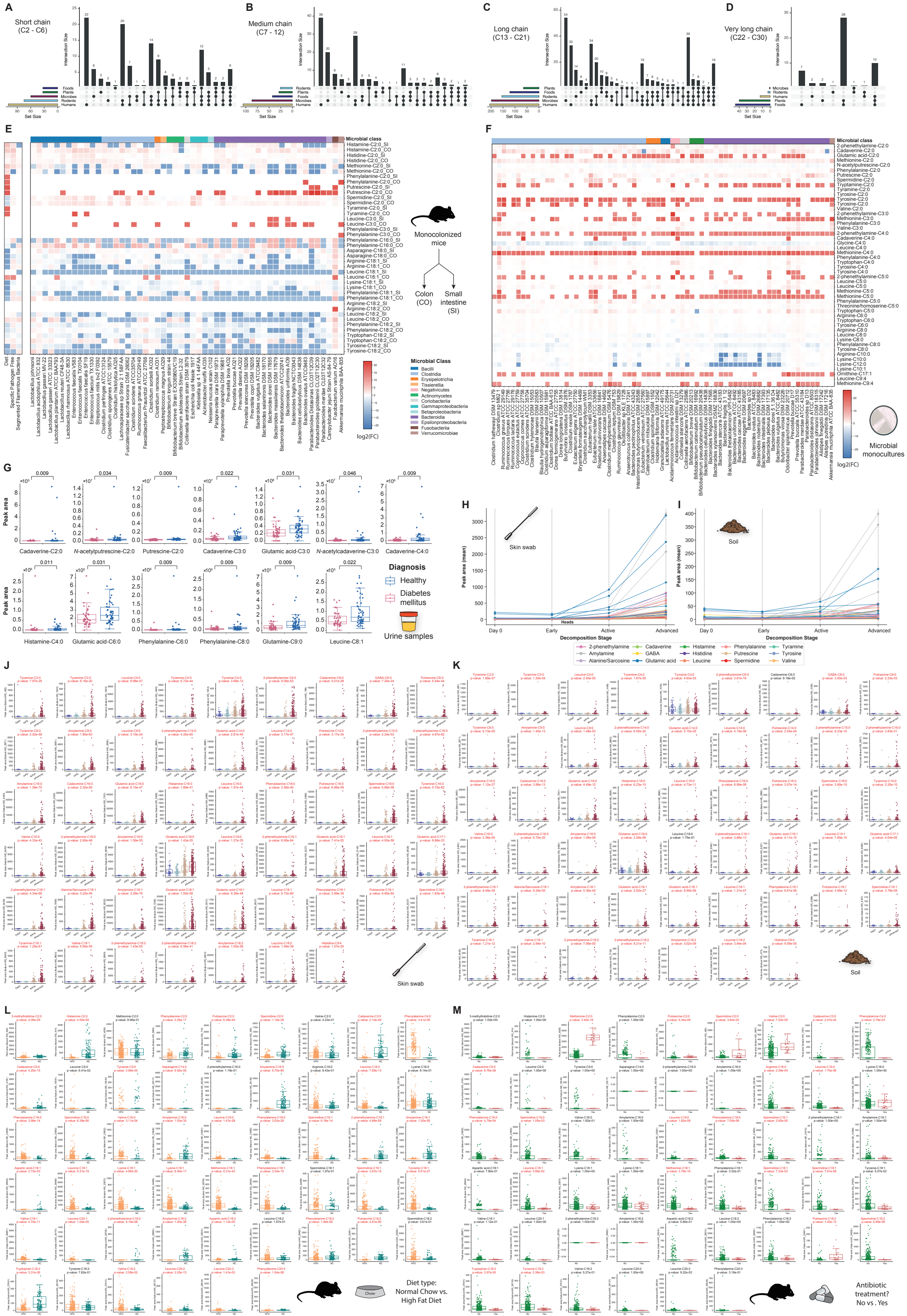


**Supplementary Figure 3. *N*-acyl lipids chain length diversity, evidence of microbial *N*-acyl lipids, and reanalysis of public datasets, related to Figure 2.** Distribution of *N*-acyl lipids in public data stratified by chain length classes. Upset plots show the number of unique *N*-acyl lipids attached to (**A**) short, (**B**) medium, (**C**) long, and (**D**) very long chain fatty acids. (**E**) Reanalysis of a public dataset of monocolonized GF mice (GNPS/MassIVE: MSV000088040, deposited in 2021)[1,2]. Heatmap log2 fold changes (FCs) of the *N*-acyl lipids matches in colon and small intestine samples of monocolonized mice relative to germ-free (GF) mice. Values of the diet, Specific Pathogen Free (SPF) mice, and of mice colonized with Segmented Filamentous Bacteria (SFB) are also shown. Red cells indicate compounds that are increasing relative to GF, while blue cells indicate compounds that are decreasing relative to GF mice. The x-axis is taxonomically ordered according to the NCBI Taxonomy ID. (**F**) Heatmap showing the log2 fold change of N-acyl lipids matches in microbial monocultures of gut commensal microbes relative to the culture media. Red cells indicate compounds that are increasing, while blue cells indicate compounds that are decreasing relative to the media. The x-axis is taxonomically ordered according to the NCBI Taxonomy ID. (**G**) Peak area abundances of *N*-acyl lipids annotated in a public dataset (GNPS/MassIVE: MSV000082261) from urine samples across clinical groups of healthy and type I diabetes mellitus. Only *N*-acyl lipids with p-values of 0.05 or less are shown. Healthy, $n$ = 52; Diabetes (type 1), $n$ = 44. (**H,I**) *N*-acyl lipids annotated from a public dataset (GNPS/MassIVE: MSV000084322, MSV000084463) of (**H**) skin swabs and (**I**) soil samples of a human cadaver decomposition study.[3] The parallel coordinates plots show the mean of the *N*-acyl lipids peak areas obtained for the different headgroups in each of the stages of decomposition. Each line represents a *N*-acyl lipid match. (**J,K**) Peak area abundances of *N*-acyl lipids annotated in public datasets from (**J**) skin (GNPS/MassIVE: MSV000084322) and **b)** soil (GNPS/MassIVE: MSV000084463) samples across different stages of decomposition of human bodies.[3] Skin: Day0, $n$ = 36; Early, $n$ = 171; Active, $n$ = 292; Advanced, $n$ = 249. Soil: Day0, $n$ = 36; Early, $n$ = 171; Active, $n$ = 299; Advanced, $n$ = 252. (**L,M**) Peak area abundances of *N*-acyl lipids annotated in a public dataset (GNPS/MassIVE: MSV000080918)[4] from mice fecal samples of mice subjected to different diets (**L**) and treatment with a cocktail of antibiotics (**M**). Antibiotics: No, $n$ = 310; Yes, $n$ = 27. Diet: HFD, $n$ = 310; NC, $n$ = 114. For the antibiotics plot, only mice fed with HFD were considered. All boxplots indicate the first (lower), median, and third (upper) quartiles, while whiskers are 1.5 times the interquartile range. Significance was tested in cases where two groups were compared using the non-parametric two-sided Mann-Whitney U test, while for more than two groups the non-parametric Kruskal-Wallis test was used, and p-values were corrected for multiple comparisons using the Benjamini-Hochberg correction. Compounds with p-values below 0.05 are highlighted in red. Icons were obtained from Bioicons.com.

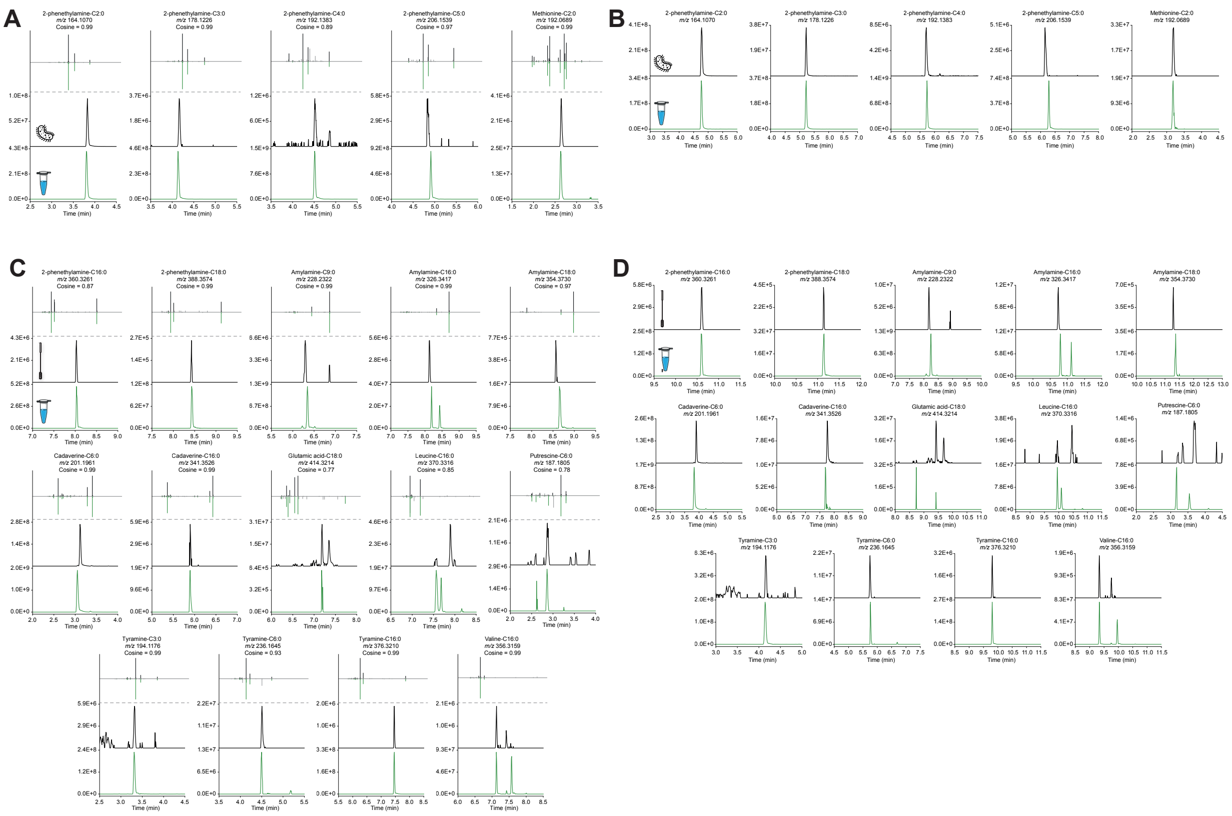


**Supplementary Figure 4. MS/MS and retention time matching of *N*-acyl lipids in samples from the microbial monocultures and from the body decomposition study, related to Figure 2.** (**A**) MS/MS mirror plots and retention time matches to *N*-acyl lipids obtained via combinatorial synthesis. MS/MS spectra on the top (black) represent spectra detected in the microbial monocultures experiment (**Supplementary Figure 3F**). An unusual series of *N*-acyl 2-phenethylamines was observed and confirmed in level 1 annotation[5,6] in two different chromatographic methods: LC1 (**A**) and LC2 (**B**) - see **Methods**. Chromatographic traces represent the exported ion chromatograms for each compound (black: sample; green: standard). (**C**) MS/MS mirror plots and retention time matches to *N*-acyl lipids obtained via combinatorial synthesis. MS/MS spectra on the top (black) represent spectra detected in the body decomposition study (**Supplementary Figure 3H-K**). Chromatographic traces represent the exported ion chromatograms for each compound (black: sample; green: standard) in two different chromatographic methods: LC1 (**C**) and LC2 (**D**) - see **Methods**. MS/MS mirror plots can be interactively inspected in the Metabolomics Spectrum Resolver[7] with the information provided in **Supplementary Table S2**.

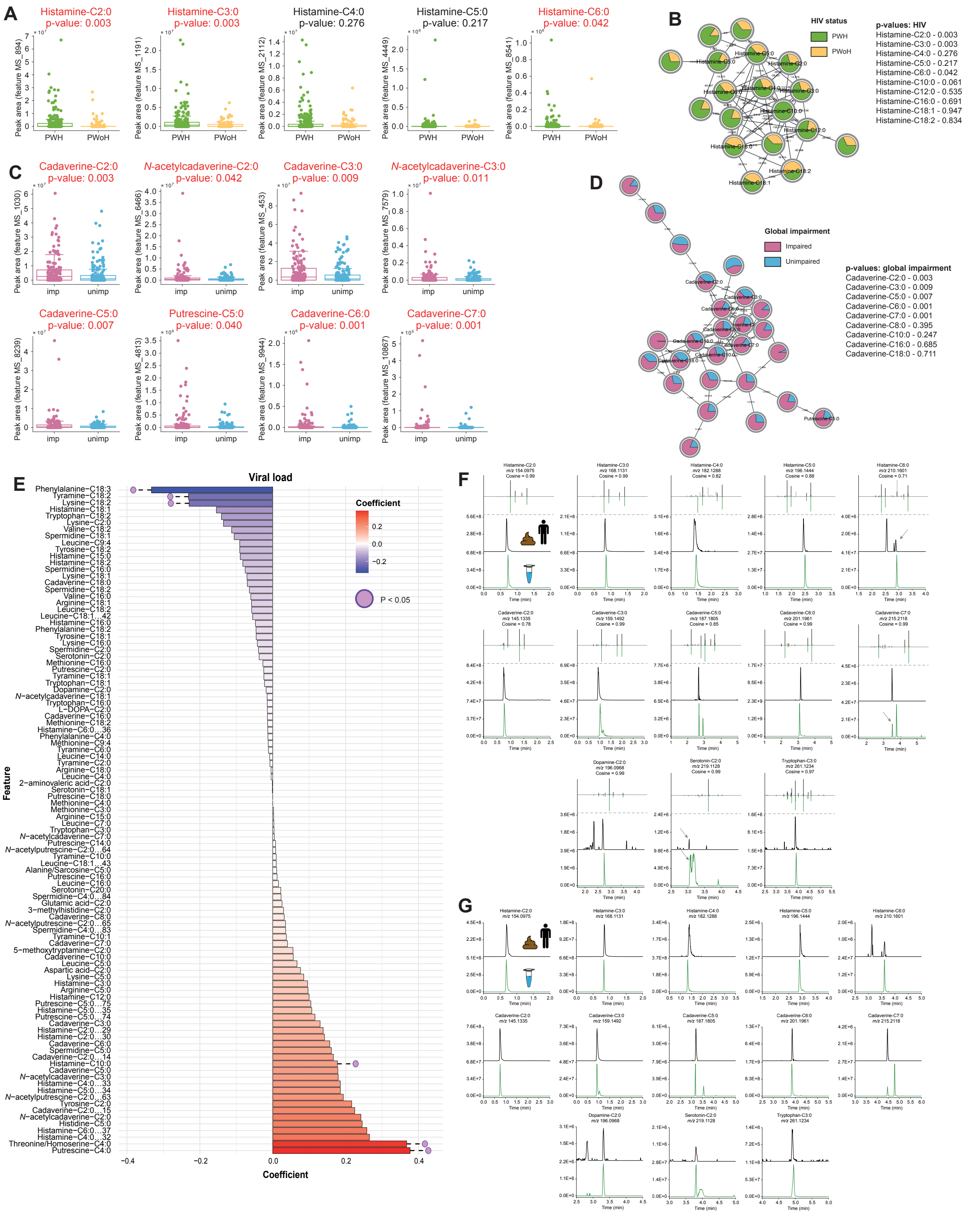


**Supplementary Figure 5. *N*-acyl lipids associated with HIV status, HIV plasma viral load, and neurocognitive impairment status, related to Figure 3.** (**A**) Peak area abundances of N-acyl histamines in people with HIV (PWH) and people without HIV (PWoH) (PWH, *n* = 228; PWoH, *n* = 93). (**B**) Molecular network obtained for histamine *N*-acyl lipids. (**C**) Peak area abundances of *N*-acyl polyamines in cognitively impaired and normal participants (impaired, $n$ = 151; unimpaired, $n$ = 162) of the HNRC. (**D**) Molecular network obtained for *N*-acyl cadaverines. Boxplots indicate the first (lower), median, and third (upper) quartiles, while whiskers are 1.5 times the interquartile range. Significance was tested using the non-parametric two-sided Mann-Whitney U test. The p-values shown are nominal p-values, and the adjusted ones (for multiple comparisons using Benjamini-Hochberg) are also available in **Supplementary Table S3**. The molecular networks were created using the Feature-Based Molecular Networking workflow[8] within the GNPS environment[9]. The nodes are annotated based on spectral similarity matches with the *N*-acyl lipids library created. The nodes represent each MS/MS spectrum, while the edges connecting them represent their spectral similarity (threshold set to cosine > 0.7). Pie charts indicate the relative abundance of ion features in each group highlighted. This dataset is publicly available in GNPS/MassIVE under the accession number MSV000092833. (**E**) Bar plots showing the correlation coefficients for the association between HIV RNA viral load and various *N*-acyl lipids in the PWH (n = 203). Red bars represent positive correlations, while blue bars represent negative correlations, as determined by linear regression models. The p-values shown are nominal; adjusted p-values (corrected for multiple comparisons using the Benjamini-Hochberg method) are available in **Supplementary Table S3**. (**F**) MS/MS mirror plots and retention time matches to the pure *N*-acyl lipids standards. MS/MS spectra on the top (black) represent the ones detected in the HNRC fecal samples, while the MS/MS on the bottom (green) are the ones obtained from the standards. Chromatographic traces represent the exported ion chromatograms for each compound (black: sample; green: standard). The chromatographic method LC1 (see **Methods**) was used. MS/MS mirror plots can be interactively inspected in the Metabolomics Spectrum Resolver[7] with the information provided in **Supplementary Table S3**. (**G**) Chromatographic traces represent the exported ion chromatograms for each compound (black: sample; green: standard), with data acquired in a different chromatographic method: LC2 (see **Methods**).

### References


1. Wu, M. *et al.* Gut complement induced by the microbiota combats pathogens and spares commensals. *Cell* **187**, 897–913.e18 (2024).
2. Song, X. *et al.* Gut microbial fatty acid isomerization modulates intraepithelial T cells. *Nature* **619**, 837–843 (2023).
3. Burcham, Z. M. *et al.* A conserved interdomain microbial network underpins cadaver decomposition despite environmental variables. *Nat Microbiol* **9**, 595–613 (2024).
4. Shalapour, S. *et al.* Inflammation-induced IgA+ cells dismantle anti-liver cancer immunity. *Nature* **551**, 340–345 (2017).
5. Sumner, L. W. *et al.* Proposed minimum reporting standards for chemical analysis Chemical Analysis Working Group (CAWG) Metabolomics Standards Initiative (MSI). *Metabolomics* **3**, 211–221 (2007).
6. Schymanski, E. L. *et al.* Identifying small molecules via high resolution mass spectrometry: communicating confidence. *Environ. Sci. Technol.* **48**, 2097–2098

(2014).
7. Bittremieux, W. *et al.* Universal MS/MS Visualization and Retrieval with the Metabolomics Spectrum Resolver Web Service. *bioRxiv* 2020.05.09.086066 (2020) doi:10.1101/2020.05.09.086066.
8. Nothias, L.-F. *et al.* Feature-based molecular networking in the GNPS analysis environment. *Nat. Methods* **17**, 905–908 (2020).
9. Wang, M. *et al.* Sharing and community curation of mass spectrometry data with Global Natural Products Social Molecular Networking. *Nat. Biotechnol.* **34**, 828–837 (2016).
